# StarCrew: A pipeline for exploratory analysis of *Starship* cargo

**DOI:** 10.64898/2026.11.19.767472

**Authors:** Andres Felipe Lizcano-Salas, Martin Ryberg, Aaron A. Vogan

## Abstract

**Background:** *Starships* are giant cargo-mobilizing mobile elements within the fungal subphylum Pezizomycotina that harbor diverse gene content, including genes for virulence and heavy metal resistance. Although *Starship* elements carry a vast array of cargo genes, the overall composition and function of this cargo remain largely uncharacterized for most elements. While existing bioinformatic tools have been designed for the identification of these elements, a gap remains in tools for characterizing their gene content while accounting for its specific characteristics.

**Results:** To address the gap in gene content analysis of *Starships*, we present StarCrew, a bash- based pipeline designed for systematically characterizing the cargo gene diversity and turnover within *Starship* elements. We demonstrate that StarCrew robustly handles datasets of varying diversity, providing the capacity to cluster elements based on collinear regions, identify nested insertion events, perform element classification, and highlight overrepresented genes within a dataset. Additionally, StarCrew serves as a diagnostic tool to flag elements requiring boundary curation. We found that the accuracy of the pipeline was sensitive to the quality of input gene models and the precision of initial element boundaries delineation. Demonstrating its utility, we identified a putative pseudokinase as a new conserved protein distributed across *Starships*.

**Conclusions:** StarCrew provides a practical and versatile resource for the continuous exploration of *Starship* gene content, designed to adapt alongside our expanding understanding of these mobile elements. Finally, we propose a complementary system to group elements with potential shared beneficial traits, based on collinear blocks, called Functional Constellations.

## 1. Background

Transposable elements (TEs) are ubiquitous DNA sequences capable of integrating into new genomic locations (1). Traditionally, TEs are classified into two classes based on their transposition intermediate: class I (retrotransposons), which use an RNA intermediate, and class II (DNA transposons), which mobilize directly via DNA (1,2). Most TEs are characterized by a simple architecture, carrying only the essential machinery required for their own mobilization in autonomous elements, or carrying only non-coding regions (1). However, recent studies in eukaryotes have identified elements that incorporate an extensive repertoire of additional genes referred to as ‘cargo’. While these genes are not required for transposition, they may confer niche-specific adaptations to the host (3). Such elements have been previously referred to as cargo-mobilizing mobile elements (CMEs) (4).

*Starships* are giant CMEs recently described as a superfamily of class II TEs in fungi. These elements are mobilized by a tyrosine recombinase (YR) that has been named captain (5,6). Primarily identified within the subphylum Pezizomycotina, these elements can reach sizes of up to 700 kb (4,5). Structurally, *Starships* are characterized by direct repeats (DRs) at their borders and, in some instances, terminal inverted repeats (TIRs) (4,5). The captain gene is situated near the 5’ border of the element, frequently starting at the first 2 kb, and is normally the first gene in the element (5,6). Most elements also harbor various cargo genes, with some hypothesized to assist in transposition activity and/or maintenance of the elements in the host, named auxiliary genes, while others are thought to confer adaptive or beneficial traits to the fungal host (4,5,7). *Starship* classification currently follows a three-level hierarchy: family, *navis*, and haplotype. The family level is defined by the phylogenetic relationships of the captain protein and is primarily identified using HMM profiles (4). Below the family level, elements are categorized into *naves* (plural of *navis*) based on the orthogroup relationships of the captain protein. Finally, at the haplotype level, elements are clustered by full nucleotide sequence similarity using a k-mer approach (4).

As previously mentioned, these CMEs carry genes that may confer selective advantages to the fungal host (7). To date, elements carrying genes for formaldehyde resistance and heavy metal resistance (including arsenic, cadmium, copper, lead, and zinc) have been characterized (6,8–10). In specific cases, such as formaldehyde resistance, experimental evidence has demonstrated that resistance is conferred by a gene cluster within these elements. Experimental deletion of the gene cluster directly correlates with the loss of the resistance phenotype, while the loss of a functional copy of a single gene from the cluster correlates with reduced resistance (8). Similar results have been obtained in experiments conducted with the *Hephaestus* element in *Paecilomyces variotii* and *Aspergillus fumigatus*, proving that the horizontal transfer of this element from *P. variotii* to *A. fumigatus* confers increased metal resistance to copper, zinc, and cadmium in the recipient strain (11). Beyond environmental stressors, elements have also been described to carry and mobilize known virulence factors in phytopathogens, such as the necrotrophic effector ToxA in *Parastagonospora nodorum*, *Pyrenophora tritici-repentis*, and *Bipolaris sorokiniana* (12,13) or the gene cluster related to the biosynthesis of *N*-lauroylethanolamine in *Verticillium dahliae* that have been showed to induce defoliation in infected plants (14,15). Furthermore, cargo genes related to pathogen survival and virulence have been described in the human pathogen *A. fumigatus* (16). Additionally, genes of relevance to the food industry have been identified within these elements, specifically those related to food adaptation, such as lactose metabolism (5,9) and a gene cluster related to salt tolerance (9). These findings showcase the critical relevance of continued study into these CMEs and their diverse cargo genes.

The current set of described elements exhibit a highly diverse repertoire of cargo genes that, in most cases, shows no similarity between elements (4). In some cases, different elements share only a small fraction of their total cargo (4,6), revealing that not only is the movement of *Starships* dynamic, but that there is also a constant internal dynamism within the element boundaries, involving the continuous gain and loss of sequences. Existing data suggests that cargo is gained or lost primarily in multi-gene blocks or gene clusters rather than isolated single genes (8,9,16,17). Currently, there is no formal description of the molecular mechanisms driving these events, nor are there bioinformatic workflows specifically designed to analyze them.

To the best of our knowledge, there are only two tools designed for *Starship*s analysis: starfish (4) and stargraph (17). However, these tools focus primarily on the systematic identification of discrete elements (4) or genomic regions potentially impacted by element insertion, termed *Starship*-like regions (SLRs) (17), leaving a gap in the characterization of cargo genes and their internal dynamics once identified. To address this, we developed the ***Star****ship* **C**a**R**go **E**xploration **W**orkflow (StarCrew), a bash-based pipeline designed to systematically characterize the cargo genes diversity and turnover within *Starship* elements (Figure 1). The primary goal of StarCrew is to provide researchers with an integrated pipeline for the exploratory analysis and comparison of cargo genes, facilitating the identification of biological patterns and the generation of testable hypotheses through bioinformatic or experimental approaches. The main two kind of analysis performed by StarCrew are Clustering analysis based on collinearity blocks and orthogroup overrepresentation analysis. Because the workflow prioritizes the identification of viable candidate genes for downstream validation, pseudogenes were excluded during workflow development. The pipeline consists of seven distinct commands and operates in a Linux environment, generating an organized project directory structure to simplify downstream analysis.

**Figure 1.**
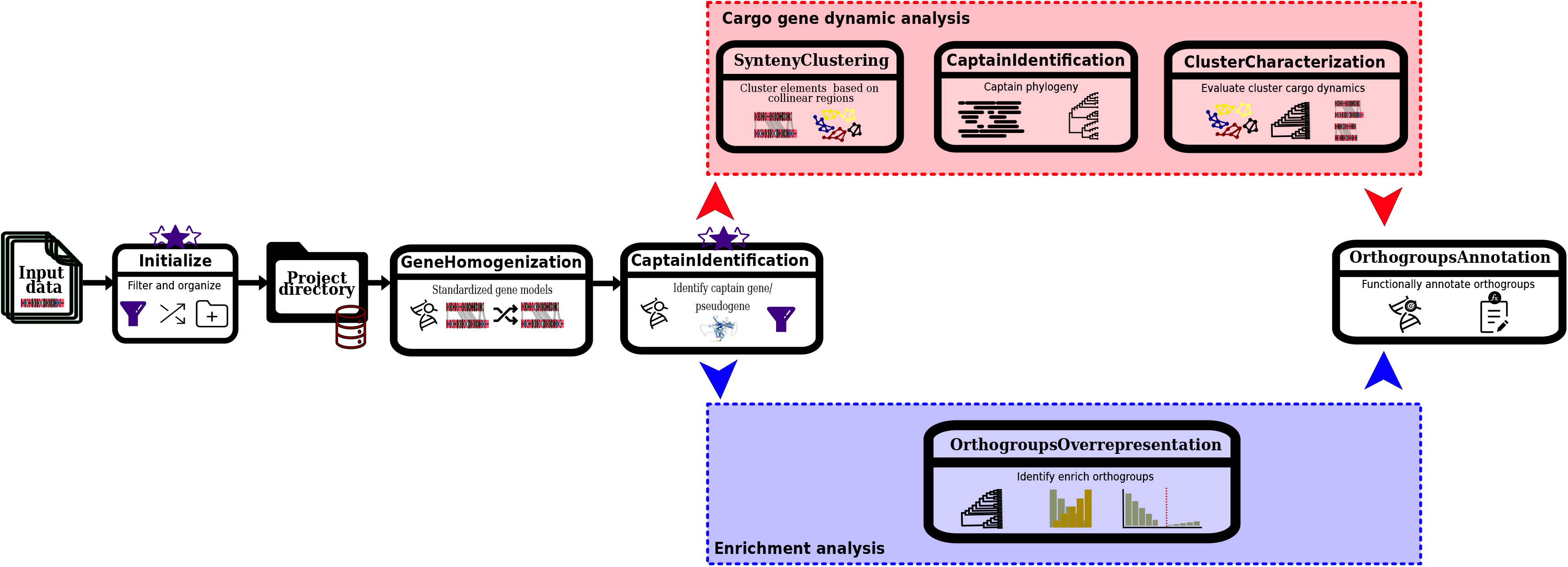
**Workflow of StarCrew**. Schematic representation of the two pipelines available in StarCrew. Each step is labeled with its corresponding command and primary objective. Purple stars denote specific stages where data filtering is performed.

## 2. Implementation

The pipeline is designed with minimal input requirements. The two primary inputs are the element nucleotide sequences in FASTA format and gene predictions in GFF3 format. Optionally, a metadata file can be supplied, containing as much information as the user requires. For more detailed information on format, and command parameters, visit the pipeline repository: https://github.com/andres2901/StarCrew.

### 2.1. Data preprocessing

The initial phase of the StarCrew pipeline involves a pre-processing step for data filtration. A mandatory minimum gene content filter excludes any elements possessing fewer genes than a specified user-defined threshold. Since the primary objective of the workflow is to analyze cargo content, retaining elements with insufficient genes is computationally inefficient, as these elements lack the gene density required to produce clustering patterns.

Furthermore, users may optionally enable filters for Repeat-Induced Point Mutation (RIP)-like signals and GC content. The optional RIP-like Signal Filtering step is designed to remove elements exhibiting extensive RIP signatures across a high proportion of their sequence; a fungal genome defense mechanism that induces cytosine to thymine transition mutations in repetitive or duplicated sequences (18–20). High RIP-like signal coverage can indicate that an element is degraded, potentially lacking functional cargo genes, or the ability to mobilize (8). For this purpose, we developed a custom Python script, inspired by theRipper web server (19), that uses three indices to define a RIP-like region: the rip substrate index, the rip product index, and the rip composite index (18–20). We modified the thresholds for the rip substrate index and rip product index based on the results presented by Margolin et al. 1988 (18) and Lewis et al. 2009 (20). It first calculates these indices across the element in 1000 bp windows with a 250 bp sliding step. A window is then designated as RIP-like only if the rip substrate index is less than 1.05, the rip product index is greater than 0.8, and the rip composite index is greater than 0. To prevent the over-identification of RIP-like regions, a RIP-like region is only established when three sequential RIP-like windows are identified, with the overlapping section among these windows defining the final region. Finally, the coverage of RIP-like regions is calculated and used to filter out elements based on a user-defined threshold.

The third filter is based on GC content; this was included to specifically remove elements that may have undergone RIP-like degradation but were not identified by the primary RIP- like signal filtering stage. This relies on the assumption that AT-rich regions are strongly associated with RIP (19,21). The allowed range for this GC content value was determined based on previous results of general GC content observed in Pezizomycotina (22,23) and prior analysis of AT-rich regions in fungi (21).

When using data from starfish (4), a preliminary process is performed prior to filtering. This stage begins with the extraction of gene information from the original genomic GFF files, which is then updated based on the specific genomic coordinates provided in the starfish metadata. Because starfish performs *de novo* prediction of captain genes (4), a modified re-prediction step is implemented to retrieve gene model details like exons, introns, and complete CDS information that are not included in starfish output. This is achieved using MetaEuk v7.bba0d80 (24) against the nucleotide sequences utilizing the YR identified by the starfish gene finder module (4) as the protein database. While starfish consistently prioritizes the *de novo* MetaEuk predicted model in cases of gene model overlap (4), this pipeline evaluates the length of all overlapping models and selects the longest available. Finally, the data is merged, organized, and subjected to the filtration steps described above.

### 2.2. Gene model Homogenization

A major challenge when integrating genomic data from multiple sources is the inherent inconsistency in gene models resulting from different prediction pipelines (25,26). These discrepancies can confound downstream comparative analyses, such as synteny mapping and orthology inference (27). To mitigate this issue, we developed a simple command designed to homogenize gene models across similar elements. Importantly, this command does not evaluate the biological accuracy of competing gene models; rather, it applies the same consensus logic used for our *de novo* captain identification by selecting the model yielding the longest protein product in overlapping regions. Consequently, the output of this homogenization step should be interpreted with caution.

To achieve this, an all-by-all BLASTN search (28) is first performed to identify element pairs sharing at least 50% coverage similarity, mirroring the BLASTN filter of synteny clustering parameters detailed below (see section “2.4. Synteny-based clustering”). Subsequently, Liftoff (29) is executed for all elements possessing at least one similar counterpart in the dataset. To optimize computational runtime, related elements are pooled into a collective "artificial genome" that serves as a composite reference against the target element. To prevent the propagation of pseudogenes, lifted gene models containing internal stop codon, and thus lacking a valid open reading frame (ORF), are filtered out. Finally, the remaining lifted genes are merged with the element’s original gene models. For any overlapping gene, the longest gene model is retained.

### 2.3. Captain Identification and phylogeny

As previously mentioned, the captain is responsible for *Starship* transposition and has been proposed as the standard for evaluating their evolution and classification (4).

Consequently, the accurate identification of the element’s captain gene is a critical task. For the purposes of this pipeline, we specifically prioritize elements for which a confident captain gene or pseudogene can be identified, while excluding those that do not meet the criteria.

To identify the captain gene, we focused on its most fundamental description: a YR containing a DUF3435 domain located in the positive strand of the 5’ region of a *Starship* (4,5). Based on this criterion, we utilize three distinct sets of HMM profiles: the captain HMM profile provided by starfish (4), the HMM profile for the DUF3435 domain (Pfam accession: PF11917.13), and a set of HMM profiles associated with the active domain of YR. This last set includes: DNA breaking-rejoining enzymes (Superfamily accession: SSF56349), Phage integrase family (Pfam accession: PF00589.27), Integrase catalytic core (CATH-Gene3D accession: G3DSA:1.10.443.10), TYR_RECOMBINASE (PROSITE accession: PS51898 version 3), and TYR_RECOMBINASE_FLP (PROSITE accession: PS51899 version 3) (30–33). For PROSITE domains, we retrieve the multiple sequence alignment from the PROSITE web database and subsequently built the HMM profiles using hmmbuild with default parameters (34).

Candidate proteins are identified by executing hmmscan (34) for each element’s proteome against these profile libraries, followed by the application of four sequential filters to define high-confidence captain genes. The first three filters are user-adjustable and include a confidence level threshold (requiring a minimum number of matches across the HMM profile libraries), a minimum amino acid length for the predicted protein, and a positional constraint. By default, the confidence level threshold is set to two, requiring matches to both the starfish captain HMM profile and the DUF3435 HMM profile, while the default minimum length is set to 250 amino acids, aligning closely with the reference YR sequence for starfish (203 aa) (4). The positional filter ensures the gene is located at the 5’ end of the element on the positive strand (or the 3’ end on the negative strand for reverse-complemented sequences) within a specified distance from the element ends. In cases where the confidence level is set below three, the pipeline prioritizes candidates with the highest number of HMM matches to ensure robust identification. The final filtering step mandates an exon count between 2 and 11; a range derived from empirical observations and existing literature. Most confirmed captains are intron-containing gene models (4,6,12,35), whereas single exon YR are frequently associated with YR-encoding retrotransposons (36). Furthermore, our preliminary analysis suggests that gene models with an excessive number of introns (exceeding 11) often represent pseudogenes. In these cases, gene predictors may introduce artificial introns to avoid in-frame stop codons, a phenomenon previously documented for other genes such as *FuSpok* in *Fusarium* (37).

For elements in which a confident captain gene cannot be identified, the pipeline performs a search for putative pseudogenes. This search is conducted using MEGABLAST (28) across both element ends, constrained by the user-defined distance parameters. The query set for this search includes captain gene CDS retrieved from the initial test dataset (Supplementary Information), specifically those containing the YR active domain, supplemented by captain gene CDS identified within the user’s current dataset. To be confirmed as a pseudogene, a candidate must span at least 600 nucleotides. Any element lacking both an identifiable captain gene and a valid pseudogene is classified as “captainless” and excluded from further analysis.

After identification of the captain gene, a phylogenetic tree is reconstructed to provide an initial characterization of the evolutionary relationships among the element captain sequences. Also, this tree serves as a reference "element tree" for subsequent orthogroup analysis. The process begins with the alignment of all identified captain genes and pseudogenes using the alignSequences subprogram of MACSE v2.07, which allows the alignment of coding sequences while accounting for premature stop codons and frameshift mutations to produce a translation-aware amino acid alignment (38). Within this pipeline, CDS from intact genes are designated as reliable sequences, while pseudogene sequences are treated as less reliable. For datasets consisting entirely of pseudogenes, a representative subset of CDS is retrieved from the pipeline database to serve as reliable reference sequences, which are subsequently removed following the alignment step. The alignment is then trimmed using Clipkit v2.7.0 (39) in gappy mode with a threshold of 0.90. The resulting alignment is then verified to determine whether the dataset contains multiple distinct sequences or is represented by identical captain proteins.

If at least two captain protein sequences are determined to be distinct, a maximum- likelihood phylogeny is reconstructed using IQ-TREE3 v3.0.1 (40). For datasets containing at least four unique sequences, the analysis employs automated model selection, allows for polytomies, and utilizes 1,000 SH-aLRT and 1,000 ultrafast bootstrap (UFBoot) iterations (-m MFP -alrt 1000 -B 1000 --polytomy) (40–43). For datasets with only two or three unique sequences, phylogenetic inference is performed without support values to ensure these data can be integrated into downstream processes, despite its lack of informative signal. Finally, Gotree v0.5.1 (44) is used to midpoint-root the tree and collapse branches that are either near-zero in length (< 10^-5^) or characterized by low support (< 80% SH-aLRT and < 95% UFBoot), when applicable.

### 2.4. Synteny-based clustering

Synteny-based clustering is performed using the computationally efficient and widely adopted MCScanX algorithm (45), integrated within the R package syntenet (46), to identify conserved collinear regions between elements. To optimize the clustering process, a preliminary evaluation of various parameters was conducted; the results of this analysis, which informed the selection of the default parameters, are detailed in the Supplementary Information. Initially, sequence local alignment is done using DIAMOND (47) in “fast” mode, employing a PAM30 scoring matrix, an e-value threshold of 1e^-5^, and a minimum query and subject coverage of 70%. Users can further refine collinearity identification by adjusting the number of required anchor points, the maximum allowed gaps, and e-value threshold. To quantify relationships between element pairs, we utilize two metrics: General Collinearity Percentage (GCP) and Element Collinearity Percentage (ECP). The GCP corresponds to the percentage of collinear genes returned by the MCScanX statistics’ output, while the ECP is calculated independently for each element (x) in a pair as 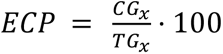, Where CGx represents the total number of non-duplicate collinear genes in element *x* and TGx is the total number of genes in that element.

Users can select returning the raw results or using one of four filtering strategies before clustering. The Strong Syntenic Pairs (SSP) strategy returns elements with a GCP greater than 41%. For elements with significant disparities in length or gene content, it requires a minimum ECP of 45% for at least one element and an 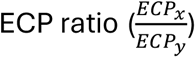 greater than 1.8 (Supplementary Information). The second strategy, the Blastn Filter, analyzes pairs not categorized as SSP via pairwise BLASTn search. These parameters were adapted from the YASS web server (48) because its accessible visualization and data summary features guided the determination of our initial parameters (Supplementary Information): -gapopen 8 -gapextend 6 -reward 5 -penalty -4 -evalue 1e^-60^. These results are filtered using a modified version of the 1.2_blastnClean.py script from Westerberg et al. (2021) (49). Individual hits failing to meet user-defined thresholds for fragment size and identity are discarded; overlapping hits are merged into contiguous regions, and regions falling below a specified size threshold are removed. Cumulative sequence coverage is then calculated to filter pairs based on a user-defined minimum coverage threshold, where at least one element should be above that threshold.

The third strategy, the Metric-Based Filter, is inspired by the ClusterBlast ranking system (50) and updates GCP and ECP values based on a scoring system. Anchor pairs are identified from the collinearity file, with the total expected score defined as twice the number of anchor pairs. Each valid hit across both DIAMOND search directions (length ≥ 100 aa; identity ≥ 60%) contributes 1 point, meaning a reciprocal hit yields 2 points. Bonus points are awarded for high-quality hits: those with ≥ 95% identity receive an additional 0.1 point for every 200 aa of alignment length. A pair is accepted if the total score meets the user-defined anchor point threshold (with a mandatory minimum of six points). Finally, the GCP and ECP are updated by multiplying them by the ratio of obtained points to expected points, capped at 1.0 to ensure values do not exceed 100%.

The fourth strategy is called Constellation, specifically designed for the grouping system designed here (see section “3.3. Functional Constellations: A Complementary system for grouping elements with share gene content”). This strategy supports two distinct filtering modes: GCP and BLASTN. In the GCP mode, all edges with weights falling below a user-defined threshold are filtered out. In the BLASTN mode, edges are filtered using the same logic as the standard BLASTN filtering strategy, with the only distinction that an edge is retained if the nucleotide alignment coverage exceeds the specified threshold for both connected elements.

Following filtration, clusters of elements with similar cargo are identified using a graph- based approach (51). A weighted undirected graph is constructed where nodes represent elements and edges represent syntenic pairs, using the GCP as the edge weight; connected components within the graph define the primary clusters. Within the Constellation strategy, an additional step is executed to split clusters based on the presence of articulation points (nodes whose removal increases the number of connected components). Specifically, when an articulation point is detected, its connections to the adjacent component exhibiting the lowest average edge weight are removed, thereby cleanly partitioning the cluster in two. To achieve a more fine-grained resolution, a subclustering process is performed on clusters containing at least four elements. An initial assessment of subclustering behavior is conducted using the Louvain method (52,53). If subclusters are detected, an iterative spectral clustering approach is employed (54,55), beginning with a k-value derived from the Louvain results. Crucially, the process only continues if this initial spectral clustering iteration produces no singletons and yields a modularity score exceeding a user-defined threshold. If these conditions are met, the process iteratively tests k+1 values until it identifies the k-value that yields the highest modularity score without generating singleton subclusters. If the initial threshold is not met, the cluster remains undivided, ensuring that subclustering is only performed when supported by significant modularity.

### 2.5. Characterization of cluster cargo dynamics

Once clusters are established, the dynamics of the elements’ cargo are explored through the detection of nested insertion events, and identification of core and accessory genes of the elements. Additionally, the pipeline evaluates potential discordances between the cargo history and the evolutionary history of the captain protein, following the assumption that the captain gene defines a *Starship* element (4). For that reason, a rooted captain phylogenetic tree must be available in a cluster to perform these steps. The only exception for this requirement occurs when conducting Functional Constellation characterization by using the “--constellation” flag (see section “3.3. Functional Constellations: A Complementary system for grouping elements with share gene content”).

The initial step involves identifying orthogroups using OrthoFinder (56) with DIAMOND (47). We employed bit-score normalization and used the captain phylogeny as the user- specified rooted “species tree”. DIAMOND parameters are adjusted to "fast" mode with a PAM30 scoring matrix, an e-value of 1e^-5^, and a minimum query and subject coverage of 70%. These modifications are intended to prioritize the identification of full-length orthologs with high identity rather than domain-level matches. Subsequently, an all- versus-all Blastn search (28,57) is performed using the same parameters as the previously described Blastn Filter strategy. When the “--constellation” flag is enabled, the pipeline additionally classifies STAR groups within each subcluster using these blastn alignment results based on pre-established coverage and identity thresholds (95% identity and 95% coverage) (see section 3.3).

Following orthogroup identification, it performs hierarchical clustering with the UPGMA method of the elements based on a similarity matrix that is created with the orthogroup gene counts, including genes not associated with any orthogroup. Subsequently, the pipeline executes an all-versus-all comparison to identify potential nesting events, defined as the insertion of one element into the sequence of another. A nesting event is identified when small elements have at least 80% of their genes belonging to orthogroups present in a larger element. The pipeline then verifies that the nucleotide sequence of the smaller element is embedded within the larger one. This verification step incorporates flexibility in the smaller element boundaries to account for potential inaccuracies in boundary definitions. Then, the command confirms that the smaller element is not simply an alignment of the beginning of the larger element; as it likely represents a truncated version of the same element or an instance of cargo gain/loss at the 3’ region rather than a discrete insertion.

Core genes are defined by a presence threshold of 80%, representing a reduction from strict (>= 99%) or soft-core (95–98%) definitions typically used in pangenomic studies (58). This lower threshold accounts for the inherent cargo variability across *Starship* haplotypes (4) and the nature of the cluster definitions. Following core gene identification, accessory genes are categorized into two categories: shared accessory and subcluster accessory. Subcluster accessory genes must meet the 80% presence threshold within a specific subcluster containing at least five elements. These subclusters are defined by the graph-based approach; however, in cases where the graph does not identify subclusters, they are defined by a height of 0.8 in the cargo hierarchical cluster tree. Any subcluster accessory gene identified in even a single element outside its group is automatically reclassified as a shared accessory gene. In “--constellation” characterization, the core gene presence threshold is increased to 90%.

Finally, the pipeline evaluates discordances between the captain’s phylogenetic history and cargo history. This comparison is intended to provide context for the distribution of core and accessory genes and is performed at the clade level. In this framework, a clade is explicitly defined as a subcluster within the hierarchical cluster tree representation. Utilizing clades rather than individual leaves allows the analysis to account for the uncertainty of bifurcations in both trees. To standardize this comparison, the captain phylogenetic tree is transformed into a hierarchical cluster tree, and the optimal number of clades is determined using the cutree() function in R (height = 0.5) and the NBClust package utilizing the "ball" index (59). The cargo hierarchical tree is then partitioned into an equivalent number of clades, enabling a direct comparison. This discordance analysis is automatically disabled during “--constellation” characterization.

### 2.6. Orthogroups overrepresentation analysis

Independent of the cargo dynamics pipeline, a secondary analysis branch was developed within the pipeline to provide an exploratory tool for identifying overrepresented or enriched orthogroups. This mode is designed to highlight potentially significant genes for subsequent validation through more robust statistical frameworks. In this pipeline, OrthoFinder (56) is executed using DIAMOND in "ultra-sensitive" mode (47), with a minimum query and subject coverage of 50%. To focus the analysis specifically on cargo, orthogroups associated with captain genes are automatically excluded. Users can select their preferred scoring matrix for DIAMOND to adjust the sensitivity for detecting orthogroups with different evolutionary distances. Overrepresentation is then evaluated through two distinct modes: Outliers and Enrichment.

In Outliers mode, the pipeline employs the Interquartile Range (IQR) fence (60,61) to identify orthogroups with an unusually high frequency relative to the dataset. Users can specify two counting methods: "Gene" or "Ship." The "Gene" method considers the cumulative number of genes within each orthogroup across the entire dataset. In contrast, the "Ship" method counts the number of individual elements containing at least one gene from a specific orthogroup, thereby ignoring paralogs to focus on element distribution frequency. After retrieving these counts, the IQR is calculated as: *IQR* = *Q*_3_ − *Q*_1_, where *Ǫ_3_* and *Ǫ_1_* represent the third and first quartiles, respectively. Given that these data exhibit a right-skewed distribution, the upper fence (UF) is identified using an adjusted approach for skewed data as follows: *UF* = *Q*3 + *ne*^3*MC*^ ∗ *IQR* (61,62), where *MC* represents the medcouple metric of skewness. Orthogroups with a size exceeding this calculated fence are categorized as outliers.

In Enrichment mode, the user selects a qualitative variable and a specific value from the element metadata for targeted analysis. To ensure a minimum level of statistical reliability, the target group must contain at least five elements. The analysis utilizes Fisher’s exact test, as implemented in the bc3net R package (63). The reference gene set encompasses all orthogroups identified by OrthoFinder, while the candidate genes are defined as those within elements associated with the selected value. Users can select their preferred method for multiple testing correction from the set available in the bc3net R package (63). Finally, an orthogroup is defined as enriched if its p-value falls below a user-defined threshold, and at least 50% of the orthogroup members are represented by genes within the specified ingroup.

### 2.7. Orthogroups functional annotation

Following the identification of orthogroups of interest from either the cargo dynamics or overrepresentation pipelines, the pipeline provides a systematic approach for functional characterization. This step operates under the primary assumption that proteins within an orthogroup maintain conserved biological functions. To ensure high-confidence functional assignments, a consensus summary is generated for any protein domain identified in at least 50% of the members within an orthogroup. The pipeline integrates three distinct annotation strategies to achieve this characterization.

First, it utilizes InterProScan (64) for a sequence-based identification of known protein domains and families. Second, it incorporates a structure-aware approach using Foldseek (65) to compare protein sequences against established 3D structural databases. In this process, protein sequences are transformed into 3Di-alphabet sequences using Protein ‘structure-sequence’ T5 (Prost5) models (66). This approach supports searches against both the Protein Data Bank (PDB) (67) and the AlphaFold database (68). Finally, HHblits (69) is employed to provide profile-level annotations for the entire orthogroup. This involves comparing a multiple sequence alignment, generated by MAFFT (70), against the Pfam-A HMM profile database (30). Ultimately, the pipeline consolidates these findings into a comprehensive summary table, integrating results from all three methodologies to provide a multi-approach functional overview of the *Starship* cargo.

### 2.8. Benchmark data

To benchmark the cargo dynamics pipeline, a dataset of 47 *Starship* elements was assembled. This collection included eight elements carrying the *Starship* formaldehyde (*ssf*) cluster (8) and 24 elements harboring arsenic resistance cluster (4,10), supplemented by 15 random elements from starbase pre-release v0.1.0 (71) (Supplementary Table S1). For the arsenic-resistant elements, inclusion was restricted to those carrying the three previously characterized core genes (9). To eliminate potential discrepancies arising from varied gene prediction methodologies in the original sources, we performed standardized gene predictions using MetaEuk (24) and BRAKER3 (72), as detailed in the Supplementary Information. The various parameter configurations evaluated are summarized in Table 1. The evaluation focused on two primary outcomes: the successful formation of clusters among elements carrying the same resistance gene clusters and the identification of these resistance-associated genes as "core" within their respective clusters.

**Table 1.** Parameter configurations for StarCrew pipeline benchmark and case study tests.

| Dataset | Pre-filter | Captain identification | Synteny-base clustering | Orthogroups overrepresentation |
|---|---|---|---|---|
| Resistance | MGC 5 | DEFAULT | 3 anchors | N/A |
|  |  |  | 3 anchors, 1e<sup>-2</sup> e-value |  |
|  |  |  | 3 anchors, 1e<sup>-2</sup> e-value, SSP filtering |  |
|  |  |  | 3 anchors, 1e<sup>-2</sup> e-value, Blastn filtering |  |
|  |  |  | 3 anchors, 1e<sup>-2</sup> e-value, no filtering |  |
| *A. fumigatus* | MGC 5 |  | DEFAULT | N/A |
|  |  |  | 3 anchors, 1e<sup>-2</sup> e-value |  |
|  |  |  | 3 anchors, 1e<sup>-2</sup> e-value, SSP filtering |  |
|  |  |  | 3 anchors, 1e<sup>-2</sup> e-value, Blastn filtering |  |
|  |  |  | 5 anchors, 1e<sup>-2</sup> e-value |  |
|  |  |  | N/A | Outlier mode, Ship count |
|  |  |  |  | Enrichment mode, navis |
| starbase | MGC 5 |  | 3 anchors, 1e<sup>-2</sup> e-value | N/A |
|  |  |  | 3 anchors, SSP filtering |  |
|  | MGC 5<br>RIP 30<br>GC 30 | CCL 3<br>length 200 | 3 anchors, 1e<sup>-2</sup> e-value |  |
|  |  |  | 3 anchors, SSP filtering |  |

Abbreviations: MGC: Minimum gene content, RIP: Repeat-Induced point mutation coverage threshold, GC: GC content threshold, CCL: Captain confidence level, SSP: Strong Synteny Pair, N/A: not applicable.

To evaluate our synteny-based approach against alternative k-mer and alignment-based strategies, we benchmarked its capacity to group elements harboring the same resistance gene clusters against sourmash v4.9.4 (73) and MMseqs2 v18.8cc5c (74). For sourmash, we evaluated three k-mer sizes (*k* = 21, 31, and 51) using a scaling factor of 100, assessing both with and without abundance weighting for the signature. Hierarchical clustering was generated using default settings in the sourmash plot command, and the resulting clusters were visually verified on the dendrogram. For MMseqs2, we executed the easy-cluster workflow using coverage mode 5 (--cov-mode 5, which calculates coverage relative to the shorter sequence) across three coverage thresholds (0.1, 0.2, and 0.4) selected to mirror StarCrew’s metric filter, blastn filter, and SSP filter logic, respectively. Additionally, we evaluated cluster mode 1 (connected component) and cluster mode 0 (greedy set cover) which partially aligns with StarCrew’s cluster and subcluster definition.

A second benchmark dataset consisted of the *Aspergillus fumigatus* starfish run from Gluck-Thaler et al. (2024) (16). For this analysis, we selected only the high-confidence haplotypes as defined in the original study, comprising 952 elements. The specific parameters evaluated for this dataset are also described in Table 1. The evaluation focused on the ability of the graph-based clustering approach to recapitulate established haplotype conformations as either clusters or subclusters.

As previous studies have not conducted any analyses of orthogroup overrepresentation, we currently lack a dataset with appropriate characteristics for testing and benchmarking. However, we emphasize that this analysis was developed as an exploratory approach to highlight potentially interesting orthogroups; consequently, any result should be paired with more robust downstream analyses.

The graph-based structures generated were analyzed using a custom Python package graph-structure v0.0.1 (available at: https://github.com/andres2901/graph_structure) that is based on NetworkX v3.6.1 package (53). For the spatial representation and visual evaluation of element clusters, networks were visualized using the Cytoscape desktop environment v3.10.4 (75).

### 2.9. Case study 1: *Aspergillus fumigatus* high confidence *Starships* dataset

Although the *A. fumigatus* dataset (16) was initially utilized for benchmarking, incongruencies were identified in its classification. For that reason, we decided to design a complementary system that can guide grouping elements with similar functions (see section “3.3. Functional Constellations: A Complementary system for grouping elements with share gene content”) and applied these criteria to create a set of these groups from the 20 high-confidence *A. fumigatus* haplotypes. The available dataset includes sequences generated by the starfish extend command (4,16) (which identify putative *Starships* regions that lack an identifiable insertion site) which in this study will be considered as “degraded” elements. Initially, a subset of these elements was defined by only maintaining non-degraded elements. Then, StarCrew was executed with this subset using the gene model homogenization and synteny clustering using the Filter metric strategy with three anchor points and e-value of 1x10^-2^. Subsequently, elements that were associated in any cluster were grouped based on their *navis* classification (done to simplify downstream analysis due to the observation of only weak connections between elements belonging to different *naves*). These *navis*-specific subsets were then processed using the Constellation strategy during the synteny clustering step, applying parameters of 4 anchor points, a maximum gap of 8 genes, and an e-value threshold of 1 × 10⁻². This analysis was executed in two parallel tracks: one with the step of gene homogenization, and another without homogenization. Both results were manually checked, and the definition of Functional Constellation was done based on (sub)cluster definition in both tracks. Elements that failed to group with any newly established Functional Constellation, and were thus considered singletons, were manually inspected to confirm true divergence. In cases where their singleton status was clearly attributable to technical issues, these elements were integrated into their appropriate Functional Constellation. Finally, to associate degraded elements with newly defined Functional Constellation, a local BLAST database was constructed using all previously excluded sequences (degraded elements). A BLASTN search mirroring our synteny clustering filter strategy was executed using an 80% coverage threshold, and the complete StarCrew pipeline was re-run with Constellation strategy. The association of each “degraded” element was done by manually checking the generated network and the characterization of (sub)clusters if available. Elements that were not included in the network were processed independently via manual curation using YASS web server (48).

Ultimately, this set of *A. fumigatus* elements also served as a case study to demonstrate the utility of the orthogroup overrepresentation module. For this specific functional evaluation, only the non-degraded element subset was utilized, and any element with more than 5% of their sequence composed of degenerate bases (Ns) were excluded. The parameters employed for this analysis are detailed in Table 1.

### 2.10. Case study 2: starbase pre-release

While the previous study case focused on the diversity of elements within a single species, we further evaluated the pipeline’s utility across a more diverse dataset using the dereplicated element set from starbase v0.3.0-pre (71). Preliminary analyses revealed inaccuracies in the taxonomic metadata associated with certain individual ship accession tags (SSB), which were subsequently inspected and corrected. While minor, undetected metadata discrepancies may remain, we are confident they do not compromise the core objectives of this study, as these inconsistencies are likely confined to the species level. Consequently, we restricted all downstream taxonomic analyses to the genus rank or higher. Dataset inclusion was restricted to elements with available taxonomy metadata. For Starbase group accession tags (SSA) containing multiple representative elements, the designated type sequence was selected; for SSAs lacking a designated type sequence, the longest element was chosen as the reference sequence. To ensure consistency in gene content, we applied the same standardized gene prediction methodology used for the resistance benchmark dataset (Supplementary Information).

The first evaluation targeted the cargo gene dynamics pipeline; the various parameter sets utilized for this testing are detailed in Table 1. Additionally, we applied our newly designed system (see section 3.3, "Functional Constellations: A complementary framework for grouping elements with shared gene content") to define preliminary Functional Constellations across elements cataloged in starbase. For this step, we selected results from the most stringent preprocessing configuration to restrict analysis to elements containing a high-confidence captain gene, excluding pseudogenized captains. Within this subset, gene model homogenization was integrated into the pipeline, and SyntenyClustering was executed in Constellation mode. Subsequently, the resulting (sub)clusters were manually inspected, problematic elements were filtered out, and a second iteration of StarCrew was executed to establish a set of preliminary Functional Constellations. A representative Constellation was then selected for downstream analysis (detailed methods provided in the Supplementary Information). For the second evaluation, using the same element subset and integrated gene model homogenization workflow, we conducted orthogroup overrepresentation analysis using Outliers mode with Ship counting. This approach allowed us to identify previously reported auxiliary genes (5,9) as well as screen for additional conserved genes. This analysis was also performed on a smaller subset excluding elements from the genus *Aspergillus*.

To confirm that the applied filters do not disproportionately remove specific clades of elements, but rather eliminate low-quality elements, we performed a phylogenetic analysis of the identified captain genes. This analysis was restricted to captain genes (pseudogenes were excluded) across three filter configurations: 1) minimum gene content of five with default captain identification; 2) minimum gene content of five, a RIP- like signal threshold of 30%, a GC content of 30%, a confidence level of three, and a minimum protein length of 200 aa; and 3) the same criteria as configuration 2, but with the captain identification confidence level reduced to two. Proteins within this dataset were classified into *Starship* families using the previously described approach based on HMMs profiles (4). Protein sequences were aligned using MAFFT v7.526 with the E-INS-i method and a maximum of 1000 iterations (70). Alignments were trimmed using Clipkit v2.7.0 in "gappy" mode with a 0.90 threshold (39). Maximum-likelihood phylogenies were then inferred using IQ-TREE3 v3.0.1 with ModelFinder, allowing for polytomies and utilizing 1,000 SH-aLRT and 1,000 ultrafast bootstrap (UFBoot) iterations (40–42). Finally, Gotree v0.5.1 (44) was used to collapse branches with near-zero lengh (< 10^-5^), and the resulting phylogenetic tree was subsequently rooted at a position facilitating comparison with the previously established *Starship* captain tree (4).

## 3. Results and Discussion

### 3.1. Collinearity-based clustering effectively groups elements by resistance gene cluster

We assembled a dataset with 47 elements to benchmark the capability of the pipeline to group elements according to the presence of two known resistance gene clusters. Five distinct parameter configurations were tested (Table 1). During initial pre-processing, four elements were excluded as no high-confidence captain gene or pseudogene could be identified. Notably, all excluded elements originated from the uncurated starbase set (71). A manual revision of these exclusions revealed that three of the four elements contained high-confidence YR but were rejected due to boundary delineation errors. In these instances, the captain genes were located beyond the first 10 kb of the element sequence; significantly exceeding the standard <2 kb range typically reported for captain gene positioning (5,6). In one extreme case, the putative YR was located in the center of the sequence. The final element was excluded due to protein length restrictions; the identified sequence was only 183 aa and lacked a functional YR recombinase domain, likely representing a non-functional gene fragment. While this specific case demonstrates that the pseudogene identification module may not be sensitive enough to retain such truncated fragments, this is a deliberate design choice. The current threshold requires a minimum of 600 nucleotides (simulating a 200 aa threshold) to identify a potential captain’s pseudogene. While lowering this length requirement might retain highly fragmented pseudogene sequences, it would likely introduce excessive noise into the captain’s gene phylogeny (76). Ultimately, these results suggest that the removal of these elements is mostly attributable to the low quality of specific uncurated sequences rather than pipeline insensitivity.

The number of identified clusters ranged from three to five across the tested configurations (Figure 2A–B; Supplementary Figure S1A-E). As expected, higher stringency resulted in a greater number of discrete clusters. For these relatively small gene clusters, the e-value used for collinear region detection proved to be a critical parameter. A comparison of two runs differing only in this value (Supplementary Figure S1B and E) demonstrates that a more stringent threshold can fragment a single cluster into two or cause specific elements to be excluded entirely. This behavior is expected because the e-value calculation is inherently dependent on the length of the sequences being compared (45). Because shorter sequences or regions are statistically less likely to achieve the low e-values typical of large-scale chromosomal alignments, using too stringent thresholds can lead to false negatives. Consequently, when the objective is to identify small regions of collinearity, we recommend utilizing more permissive e-values to maintain cluster integrity. Additionally, our comparison of filtering strategies shows that metric-based filtering is the most inclusive (least stringent) approach, whereas the SSP filtering strategy is the most conservative.

**Figure 2.**
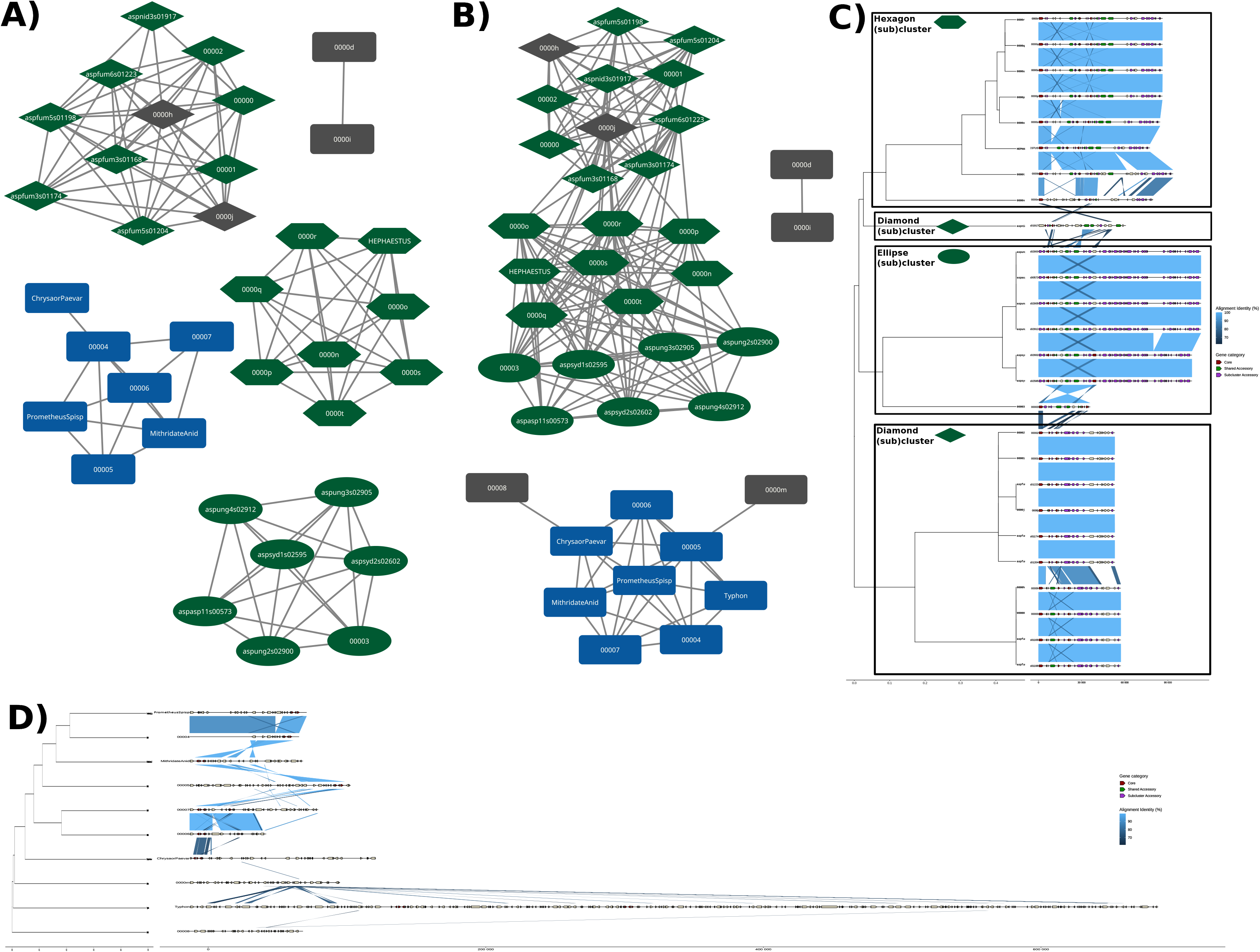
**StarCrew Benchmark of cargo dynamic analysis using the resistance gene cluster dataset**. Graph representations using Prefuse force direct layout of syntenic- based clustering across the benchmark dataset with three anchor points and **A)** e-value of 1e^-2^ and Blastn filtering strategy, and **B)** e-value of 1e^-2^ without filtering, Node shapes denote (sub)cluster divisions. Node colors indicate the origin set: green (arsenic resistance), blue (*ssf* set), and gray (random starbase elements). Synteny characterization analysis of the **C)** arsenic cluster and **D)** formaldehyde cluster (modified for clarity from Supplementary Figure S1G), derived from the three anchor points, e-value of 1e^-2^ with metric-based filtering test. Rectangles in Panel C indicate which (sub)cluster from panels A and B is represented by each clade.

Regarding the two resistance clusters of interest, elements harboring arsenic resistance genes were grouped into one to three clusters depending on the parameters used. Interestingly, the three clusters generated under the most stringent test corresponded to the three subclusters identified in the less stringent configurations. This consistency highlights the reliability of the subclustering algorithm in identifying relevant divisions within a connected component. Our analysis suggests these divisions are driven by taxonomic relationships and structural variations between related species. For example, one subcluster consists primarily of *Aspergillus fumigatus* elements and is composed of copies from two distinct elements (Diamond (sub)cluster in Figure 2C, Supplementary Figure S1F). In contrast, the non-*fumigati Aspergillus* subcluster is dominated by an element from *Aspergillus sydowii* that is nested within other elements from related species (Ellipse (sub)cluster in Figure 2C, Supplementary Figure S1F). Importantly, despite these subdivisions, the three previously characterized core genes of the arsenic resistance cluster (9) were successfully identified as "core" across all tested configurations, showing the reliability of the core gene identification.

The *ssf* cluster, associated with formaldehyde resistance, demonstrated a more robust clustering pattern, with no fragmentation observed between associated elements across most configurations (Figure 2A-B, Supplementary Figure S1A-E). Only the most stringent parameter set resulted in the exclusion of two elements; this was attributed to their limited syntenic connection to the rest of the group, which was restricted exclusively to the resistance gene cluster itself. Additionally, two elements that do not harbor the *ssf* cluster were grouped within this cluster (Figure 2D, Supplementary Figure S1G). This result is expected given that the pipeline identifies synteny in an unbiased manner; any shared collinear region, regardless of specific gene function, will drive clustering.

Within the *Typhon* element, we successfully identified the duplication of the *ssf* cluster as previously reported (8). However, the preliminary analysis only partially recovered the six previously reported core genes (8). Within the initial cluster of eight elements, only four genes (*ssfA*, *ssfB*, *ssfC*, and *ssfX*) met the core threshold. When the *Typhon* element was filtered under more stringent parameters, the core gene count increased to six (Supplementary Figure S2). Five of these correspond to the previously characterized *ssf* genes (adding *ssfF*), while the sixth is a putative *uvrA* homolog, a member of the bacterial *uvrABC* DNA damage recognition complex (77,78). Detailed inspection of the domain architecture revealed that the homolog present in these elements retains the DNA- binding domain but lacks the *UvrB*-interaction domain (77). This suggests it may maintain a role in DNA damage recognition; notably, no other components of the *uvrABC* system have been described in fungi (6,8). This *uvrA* homolog appears to be mobilized near the *ssf* cluster in several elements (Supplementary Figure S2), a close relation also noted in prior studies (6,8). Interestingly, the experimental characterization of the *ssf* cluster was conducted using the *Chrysaor* element, which lacks this gene (8), confirming that *uvrA* is not required for formaldehyde resistance. Nevertheless, its consistent physical linkage to the *ssf* cluster in otherwise distantly related *Starships* is intriguing and warrants further investigation into its potential functional role within these elements.

The discrepancies between our core gene identification and previous report are primarily associated with gene prediction challenges rather than StarCrew cluster characterization failure. Because gene prediction is an external prerequisite that users must perform prior to running the workflow. For instance, the *ssfD* gene model is missing in the *Chrysaor* element from *P. variotii*, and *ssfF* is absent in both the *Mithridate* element of *Penicillium chrysogenum* and *Aspergillus subversicolor* IBT 32280. These missing gene models directly explain why these specific genes did not meet the threshold to be classified as ‘core’. Furthermore, under these circumstances, the gene homogenization step will not improve core identification outcomes due to the low sequence similarity among all elements harboring this gene cluster; for this reason, this step was omitted from this benchmark analysis.

Comparing our synteny-based clustering against MMseqs2 and sourmash reveals that sequence-alignment and k-mer approaches fail to group elements according to their shared gene cluster (Figure 3). For MMseqs2, connected component clustering merged most elements into a single cluster (Figure 3B), whereas greedy set cover generated clusters that intermixed elements carrying the arsenic resistance cluster with those carrying the *ssf* cluster (Figure 3C). In the case of sourmash, default hierarchical clustering failed to consolidate elements harboring the arsenic cluster into a single group, instead scattering them across the four annotated clusters (Figure 3C). Notably, the similarity matrix reported zero k-mer similarity between two distinct clusters that both contain elements carrying the arsenic resistance gene cluster (Figure 3C, orange vs green cluster). These results demonstrate that synteny-based clustering provides a distinct advantage when grouping elements by shared functional potential compared to nucleotide-based methods. While nucleotide sequence similarity across varying thresholds does not guarantee shared functional roles, synteny-based analysis focuses specifically on conserved gene suites that can guide functional interpretation.

**Figure 3.**
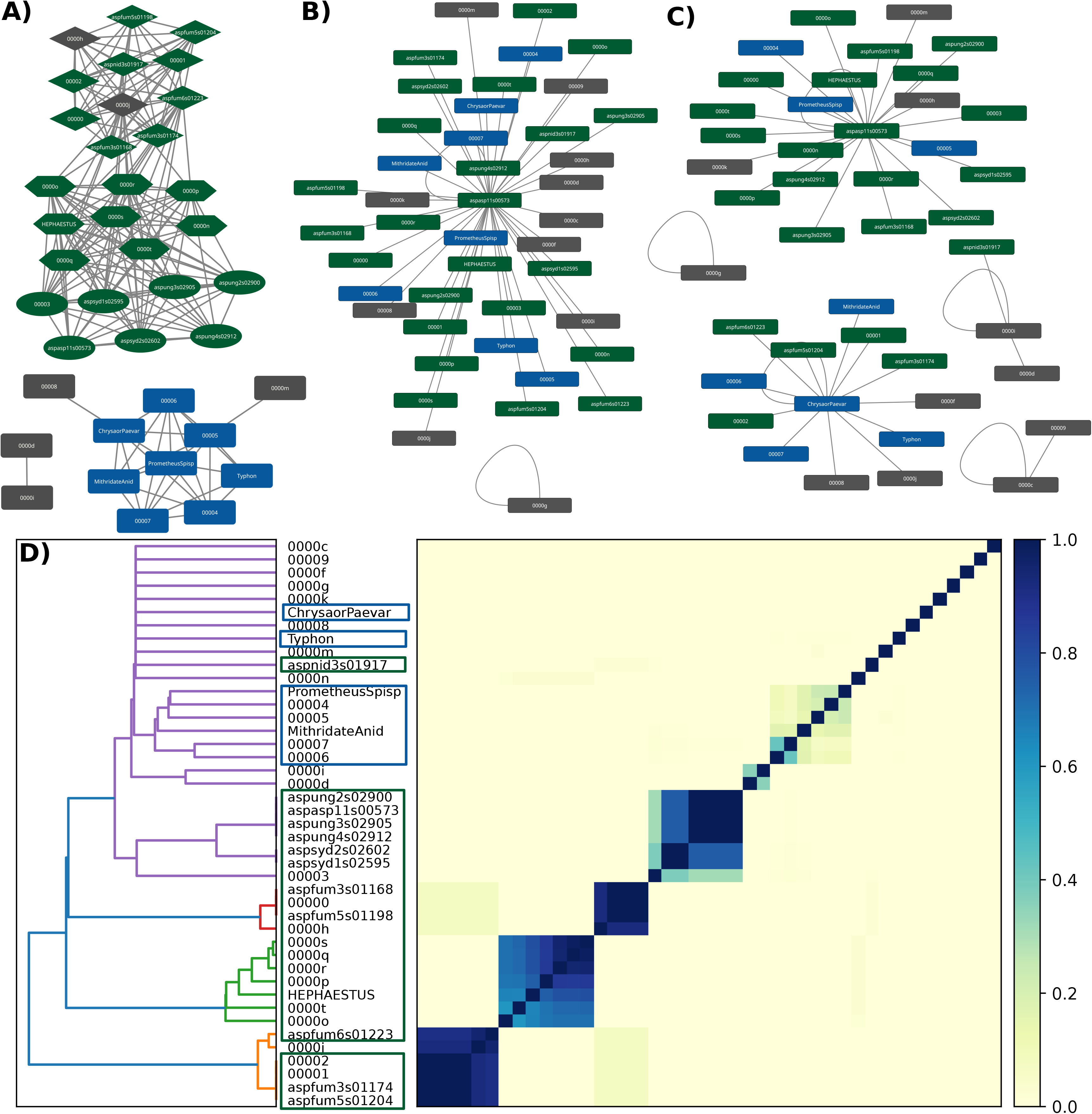
Benchmark comparison of *StarCrew* synteny-based clustering against alternative methods using a resistance gene cluster dataset. A) Graph representation of *StarCrew* synteny-based clustering across the benchmark dataset generated using three anchor points and an *E*-value threshold of 10⁻² without filtering. Node shapes denote assigned subcluster divisions. **B)** Network visualization of MMseqs2 clustering output generated using a coverage threshold of 0.1 and connected component cluster mode. C) Network visualization of MMseqs2 clustering output generated using a coverage threshold of 0.1 and greedy set cover cluster mode. **D)** Similarity matrix generated by *sourmash* using a *k*-mer size of 51 with signature abundance weighting; clade colors on the dendrogram correspond to automatically assigned hierarchical clusters. Node colors in **(A), (B)** and **(C)**, and perimeter rectangle annotations in **(D)** denote the dataset of origin: green (arsenic resistance), blue (*ssf* set), and gray (random starbase elements).

The demonstrated efficacy of StarCrew in clustering elements based on a single shared gene cluster highlights its ability to identify collinear regions and group elements based on these conserved segments which could help into identifying elements with share beneficial trait as suggested for those carrying the arsenic resistance cluster (10) or *ssf* cluster (8). However, as evidenced by the *ssf* cluster, the sensitivity of core gene identification is heavily reliant on the quality of the underlying gene models. Inconsistent gene predictions can introduce significant noise, potentially lowering a gene’s presence proportion below defined thresholds and masking legitimate core components. This underscores that while the pipeline is generally robust, its output is limited by the quality of the input annotations. Consequently, we recommend users to evaluate their data for fragmented or missing gene models. Such technical artifacts can directly obscure the identification of true core cargo. Furthermore, as previously noted, elements excluded during the Captain Identification phase should be manually inspected; these cases might represent biologically valid elements that simply require boundary curation to correct for inaccurate element delineation.

### 3.2. The Synteny-based graph highly correlates with haplotype definition in *Aspergillus fumigatus* dataset

To evaluate StarCrew in a scenario representative of large-scale intra-species pangenomic studies, we analyzed 952 high-confidence *Starship* elements from *A. fumigatus* (16). Across five distinct parameter configurations (Table 1), the network assortativity for established haplotypes remained exceptionally high, ranging from 0.987 to 0.998. This indicates that elements connect almost exclusively with members of their own haplotype, confirming the robust discriminatory power of our synteny-based clustering (Figure 4, Supplementary Figure S3).

**Figure 4.**
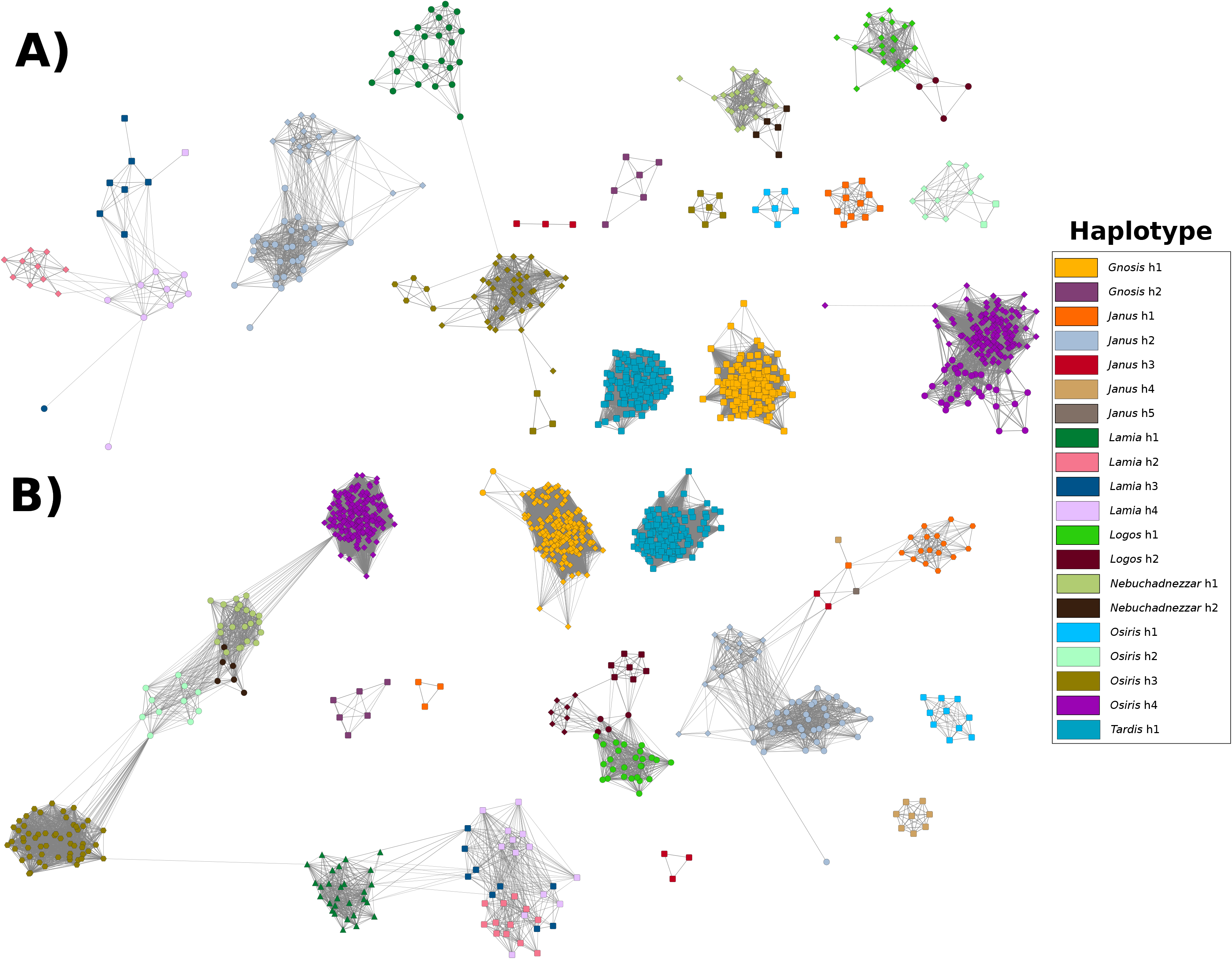
**StarCrew Benchmark of cargo dynamic analysis with *Aspergillus fumigatus* dataset**. Graph representations of syntenic-based clustering across the benchmark dataset with **A)** default parameters, and **B)** three anchor points an e-value of 1e^-2^ and metric-based filtering strategy. Node color denotes previous haplotype classification, while node shape indicates the subcluster classification assigned by StarCrew. Edges represent the general collinearity percentage (GCP), with a Prefuse force-directed layout applied to position nodes according to GCP weights.

While many haplotypes remained isolated across all tests (*Gnosis* h1/h2, *Tardis* h1, and *Osiris* h1), others exhibited inter-haplotype connectivity (i.e., *Lamia* h2/h3/h4, *Logos* h1/h2, and *Nebuchadnezzar* h1/h2) or were further separated in subcluster by the pipeline. For instance, the subclustering of *Osiris* h3 was driven mostly by significant size variation (Supplementary Figure S4). In contrast, the subclustering of *Logos* h2 was caused by technical discrepancies in gene prediction; one subcluster contained "broken" or truncated gene models (Supplementary Figure S5A) compared to the gene models in the corresponding position on the other subcluster (Supplementary Figure S5B).

StarCrew revealed significant discordances in haplotype classification within the *Janus navis* (Supplementary Figure S6), which was originally described as containing five distinct haplotypes in *A. fumigatus* (16). Notably, *Janus* h1 was split into two separate clusters even under the least stringent parameters due to lack of collinearity. This structural divergence was confirmed by synteny analysis between the three elements in the smaller cluster and two elements in the larger *Janus* h1 cluster (Supplementary Figure S6B). Interestingly, all elements in the smaller *Janus* h1 cluster possess pseudogenized captain genes, suggesting these elements are no longer active. By analyzing the flanking contig sequences, we confirmed that these elements reside at the exact same genomic location. Furthermore, cross-referencing these isolates with the species-wide population structure of *A. fumigatus* (79) reveals that the strains harboring this inactive variant are highly close phylogenetically, pointing to a single *Starship*- retention event. We also identified a "bridge" subcluster composed of elements from *Janus* h1, h3, h4, and h5, which links the h1 and h2 haplotypes (Supplementary Figure S6A and S6C). Furthermore, *Janus* h2 was divided into two subclusters, likely due to a high frequency of structural variants (Supplementary Figure S6D). Within this haplotype, one element was connected only by a single edge in the two less-stringent parameter sets (Figure S6B, E) and was entirely absent from the other three tests. Synteny analysis confirmed that this tenuous connection is primarily driven by a gene cluster at the terminus of the element containing small, likely fragmented, gene models that are not conserved across other members of the *Janus* h2 haplotype (Supplementary Figure S6E). Further review highlighted additional inconsistencies in the *Janus* h2 group, where two elements exhibited synteny only over the first ∼12 kb (Supplementary Figure S6E). This shared region represents only three genes (including the captain) and accounts for a mere 28.84% of the smaller element. Under the current definition of *Starship* haplotypes (4), such limited homology is insufficient to justify classifying these elements together. Given that the value of *Starship* haplotype classification relies on defining a cohesive cohort of element copies with shared cargo (4), grouping elements with near-zero gene conservation under a single haplotype misleads downstream analysis.

In the *Lamia navis*, other sets of discordances arise, in this case between elements of haplotypes h2 to h4 (Figure 4; Supplementary Figure S3, S7A-C). One clear example is a set of six elements that have been previously classified to belong to *Lamia* h3 and h4 (Supplementary Figure S7C). The misclassification of two of these elements is confirmed as in the test with default parameters these elements were grouped with the other haplotype rather than the one they were initially classified (Supplementary Figure S7A). The synteny plot from this subcluster shows a good level of synteny between these six elements (Supplementary Figure S7D). Another major difference between these six elements and the rest of the elements in their respective initial haplotypes is the size; these elements are less than half the size compared to elements in their original haplotype. A second classification discrepancy involves an element assigned to haplotype h4 despite its graph topology placing it closer to haplotype h2 (Supplementary Figure S7B, C). Synteny analysis indicates that while high nucleotide sequence identity is maintained, differences in gene models between the haplotypes likely contributed to the discrepancy. However, the most significant discriminatory factor is again element size: the discordant element exhibits a size profile much more characteristic of *Lamia* h2 elements than the significantly larger elements typical of *Lamia* h4 (Supplementary Figure S7E).

Given the issues identified in these two *naves*, we evaluated how well connected all haplotypes were across each parameter configuration (Supplementary Table S2). Generally, for the remaining haplotypes, the majority of connectivity issues were found to stem from discordant gene models rather than a lack of collinearity. However, as observed in *Osiris* h3, specific subclustering events in other haplotypes (such as *Osiris* h2) were driven by variation in element size. This size division was also observable in the edge weight distribution; even when these haplotypes maintained high density, they exhibited a relatively low mean edge weight compared to other well-defined haplotypes (Supplementary Table S2).

On the other hand, the core gene identification successfully retrieved previously reported genes with functions of interest, such as *lysM* and *GH18* in *Gnosis* h1, *GH71* in *Tardis* h1, and components of the *fumihopaside A* BGC in *Gnosis* h2 (16). However, in *Lamia* h1, the previously reported *GH18* and *LysM* genes (16) were only present in ∼25% of the elements. Because StarCrew identifies core genes based on consistent presence across the entire cluster, it correctly excludes them. This discrepancy demonstrates that relying on a single ’type-element’ can be misleading, as it fails to account for the internal heterogeneity and cargo turnover within a haplotype.

Although the *A. fumigatus* dataset was initially utilized for benchmarking, the identification of these inconsistencies underscores the utility of StarCrew for future studies. Our results demonstrate that the pipeline offers distinct advantages as a specialized tool for classification refinement. Also, this suggests that relying solely on nucleotide k-mer approaches for classification may be insufficient for certain *naves*, necessitating more robust methodologies.

### 3.3. Functional Constellations: A Complementary system for grouping elements with shared collinear blocks

Our preliminary results highlight clear advantages of clustering elements based on collinear regions. Accordingly, we propose a complementary system for grouping elements with shared functional potential, termed "Functional Constellations" (hereafter referred to simply as Constellations), based on conserved collinear blocks. This proposal operates on the premise that *Starships* that consistently maintain a high proportion of collinear blocks may confer a common beneficial trait. Consequently, defining a Constellation requires identifying the core cargo gene suite that unites all constituent elements. We consider that this perspective effectively accounts for cargo genes’ turnover when studying the host-adaptive potential of these elements. By analogy to astronomical constellations composed of stars, a Constellation is built from ***ST****arship* **AR**rays (STARs), defined here as *Starship* copies sharing at least 95% nucleotide identity across 95% sequence coverage. These thresholds align with those established in starbase for defining SSAs (71). Before establishing Functional Constellations, researchers should systematically address five questions:

**1. What is the minimum gene content required for an element to be included in a Constellation?** Because Constellations are thought to communicate likely shared functional or ecological advantages to the host, elements lacking substantial cargo genes provide limited comparative utility. While StarCrew utilizes a default minimum threshold of five genes, this baseline parameter should ideally scale upward for datasets characterized by elements with exceptionally high gene density.
**2. What constitutes the minimum core gene set defining a Constellation?** To communicate a cohesive beneficial trait, all elements assigned to a single Constellation should share a set of core cargo genes, explicitly excluding the captain gene. We propose starting with a minimum of four core genes. These core genes do not need to be strictly unique to a single Constellation; overlapping profiles may occur, particularly between nested or structurally expanded copies.
**3. What is the maximum permissible size variation within a single a Constellation?** An expanded element can harbor accessory genes that might offer distinct traits, even if it shares an entire core gene set with a smaller counterpart; consequently, those expanded variants should be classified into a separate Constellation. As a baseline starting point, we suggest that the smallest element must encompass at least 80% of the size of the largest element. While this threshold may be flexibly calibrated between 70% and 90%, setting a threshold below 70% risks defining incongruent groups as observed in the current haplotype definition in the *A. fumigatus* dataset.
**4. Should size variation be calculated by nucleotide length or gene count?** We recommend that this metric be determined empirically by researchers based on the perceived relevance of non-coding regions within their organism of interest. If non- coding regions are considered functionally negligible, element size may be calculated purely by gene count. Conversely, if these intergenic regions are expected to harbor critical regulatory non-coding sequences, size should be calculated relative to total nucleotide length. Furthermore, selecting nucleotide length over gene count is highly recommended when automated gene prediction within these elements shows inconsistencies.
**5. What proportion of an element’s total gene content must the core set represent?** Synthesizing the concepts of size variation and functional conservation, we propose that the core gene set should account for at least 80% (with a flexible range between 70% to 90%) of an individual element’s total gene model count. A significantly lower percentage implies that the element has expanded substantially beyond the Constellation definition, potentially introducing novel, independent phenotypes.

By unifying the answer to these five questions, a standardized reference point can be established: “A *Starship* Functional Constellation is defined as a cohesive group of elements each containing a minimum of five genes that all share at least four core cargo genes with each other. These core cargo genes must constitute at least 80% of the total gene content of each element within the group, and the size difference between the largest and smallest members must not exceed 20%”. However, we consider that the specific thresholds of this definition are modifiable while staying in line with the main logic of identifying a set of elements with putative similar beneficial trait.

This system is proposed as an independent concept rather than a formal classification scheme like that previously described (4). Rather than requiring elements to belong to the same *navis* to be grouped together, this system evaluates cargo gene similarity, allowing elements across distinct *naves* to be clustered based on shared cargo content. Another crucial aspect that we include in the system is the capacity to not only group elements based on collinear blocks but also to differentiate them based on their structural or functional state. For now, because little is known about the complete life cycle of a *Starship* copy, we propose flagging two specific states of a STAR: non-mobile and degraded.

Under this system, a "non-mobile" element is defined as one that harbors a pseudogenized captain gene but retains precise structural boundaries and exhibits no other signs of functional decay. This state may suggest that pseudogenization was only recently initiated and that the cargo genes may still be fully functional, even if the element itself is no longer mobile. This transition may represent a natural phase of the *Starship* life cycle, leading toward the eventual fixation of beneficial cargo genes within the host genome (8). On the other hand, within this system, a "degraded" element is characterized by elevated signatures of Repeat-Induced Point mutation (RIP) or other genome defense mechanisms, and/or large-scale structural variations that obscure well-defined boundaries and/or reliable insertion site identification.

Some degraded elements can still be confidently assigned to a Constellation if they maintain significant nucleotide sequence conservation; specifically, sharing at least 80% alignment coverage (with a flexible range of 70% to 90%) relative to the smallest intact element within the target Constellation. An example of this scenario was previously identified in *Prometheus* family elements harboring the *ssf* cluster (8). Critically, these degraded elements cannot be used to establish the initial set of Constellations and may only be mapped onto previously defined Constellations. This requirement is thought to prevent the establishment of Constellations solely composed of degraded elements that could confound downstream functional inferences.

To establish a Constellation, the group must be represented by at least two non- degraded copies. These two representative copies may either define two distinct STARs or represent copies of the same STAR captured across different host genomes. To enable automated clustering under this system, we implemented the Constellation mode within the SyntenyClustering command of StarCrew (See section “2.4 Synteny-based clustering"), and modifications were integrated into the ClusterCharacterization command to accommodate Contellation characterization (See section “2.5 Clustering Characterization").

To standardize reporting across studies, we propose a unified nomenclature code based on species designation, Constellation assignment, STAR group, and element functional state. Intact and putatively mobile copies should follow the convention sp-code FC#S#, where sp-code represents a six-letter species identifier (the first three letters of the genus followed by the first three letters of the species name), FC# denotes the assigned Constellation number, and S# designates the STAR number (e.g., Aspfum FC1S1). Elements that have lost mobility due to captain pseudogenization but remain structurally intact are explicitly flagged to denote their status using the nomob- prefix (sp-code nomobFC#S#). Degraded elements carry an explicit deg- prefix; if a degraded copy can be unambiguously assigned to a specific STAR within a Constellation, it is designated as sp-code degFC#S#, whereas if STAR resolution remains ambiguous but the overall Constellation is clear, it is designated as sp-code degFC#.

Furthermore, when describing Constellations associated with expected beneficial traits, researchers are encouraged to focus primarily on the defined core genes of that Constellation. If a representative "type" element must be selected for downstream analysis or reference, we recommend choosing an individual element whose cargo consists entirely of the defined core genes of that Constellation. Selecting elements that carry additional, non-conserved, genes may introduce unnecessary noise which can confound comparative analysis and mislead functional interpretations of Constellation behavior.

Finally, while this system was conceptualized primarily for intra-species pangenomic analyses, its logic remains highly applicable to multi-species comparative genomics, particularly when tracking cases of horizontal element transfer across species. For multi-species contexts, the sp-code is modified to a four-letter prefix comprising the first letter of the lowest shared taxonomic rank followed by the first three letters of that rank’s name (e.g., Genus *asp* = Gasp for a Constellation shared across multiple *Aspergillus* species).

### 3.4. A set of Functional Constellations from the high- confidence haplotype dataset of *A. fumigatus*

Applying this new system, we defined a set of Constellations from the *A. fumigatus* dataset of 20 high-confidence haplotypes (16). Our primary objective was to generate a congruent set of groups to facilitate a deeper understanding of the *Starship* repertoire in *A. fumigatus*. Due to identified gene prediction discrepancies, we utilized nucleotide length as our metric for element size. Element boundaries were considered accurate for 463 elements, as the previous study indicated that manual boundary curation had already been performed to ensure high-confidence elements (16).

For the Constellation definition, 463 elements from the complete 952-element dataset were selected because they possessed clearly defined boundaries. Among these, 413 elements (∼89.20%) were automatically grouped to 24 Constellations, while 8 elements were manually added to the Aspfum FC1 and FC2 Constellations, leaving the remaining 42 elements (∼9.07%) outside any Constellation (Supplementary Table S3 and S4). Overall, although the Constellation system does not explicitly enforce a *navis* constraint, our preliminary analyses revealed that Constellations in this dataset were strictly restricted to elements belonging to the same *navis*. In this grouping system, elements from previously described haplotypes (*Janus*-h2, *Janus*-h4, *Janus*-h5, and *Lamia*-h4) were not included in any of the Constellations because their constituent elements were resolved as singletons. Furthermore, the previous haplotypes *Gnosis*-h1, *Janus*-h1, *Lamia*-h1, *Logos*-h2, *Nebuchadnezzar*-h1, *Osiris*-h3, and *Tardis*-h1 were split in two or three distinct Constellations, an outcome driven primarily by significant intra-haplotype size discrepancies among their component elements that did not align with this system.

Overall, no substantial differences in clustering were observed between workflows executed with or without the gene homogenization step. The primary distinction lays in the collinearity percentage (the proportion of genes located within a syntenic region), which increased following gene homogenization. Notably, for Constellations that contained elements from the *Osiris* and *Tardis naves*, gene homogenization yielded an increase in the number of collinear pairs; furthermore, in both cases, the step facilitated the inclusion of two additional elements that were otherwise excluded when homogenization was omitted. However, in the specific case of Aspfum FC1, the gene homogenization step introduced significant artifacts in the gene models. Consequently, for the Aspfum FC1, we utilized the clustering results generated without gene homogenization for core gene definition. This outcome directly supports our prior assertion in the "**Implementation**" section that the outputs of the gene homogenization workflow must be interpreted with caution. For the remaining Constellation, we retained the core gene definitions obtained via the gene homogenization step.

As proof of the utility of this new system for scientific communication, we highlighted the division of the previously described *Nebuchadnezzar* h1 haplotype into two distinct Constellations, designated Aspfum FC5 and FC11. A primary distinction between them is that Aspfum FC11 is slightly larger than Aspfum FC5 due to a ∼10 kb variable region near the 5′ boundary, where the cargo gene content differs between the two groups (Supplementary Figure S8). Crucially, this variable region harbors the arsenic resistance gene cluster previously identified as a function of interest within the original *Nebuchadnezzar* h1 type element (16). Therefore, this division will clarify which strains harbor this gene cluster mobilized by a *Nebuchadnezzar* element.

As proof of utility for the two-tiered hierarchy between Constellation and STAR, we identified that Aspfum FC7 comprises two distinct STARs whose primary difference involves the fumigermin biosynthetic gene cluster (Supplementary Figure S9). Specifically, Aspfum FC7S1 contains all four tailoring genes previously reported for this BGC, whereas Aspfum FC7S2 lacks three tailoring genes while retaining the core polyketide synthase (PKS) gene and oxygenase gene. Crucially, prior studies have demonstrated that these tailoring genes are not required for fumigermin secondary metabolite production (80). Establishing this finer-scale distinction within a Constellation might facilitate the communication of how this BGC has been horizontally transferred across strains compared to treating all elements as a single group.

Subsequently, the 489 "degraded" elements, which in this analysis comprised all elements identified via the starfish extend command (16), were mapped against the newly defined Constellation. In total, 145 "degraded" elements (29.65%) were successfully integrated into these Constellation, underscoring the substantial divergence in length and gene content between the degraded and non-degraded set.

In summary, the Constellation set of *A. fumigatus Starships* corresponds to 567 elements out of the original 952, representing ∼59.56% of the original dataset size. Metadata and element designations for this set are provided in Supplementary Table S3. In retrospect, a primary issue limiting the inclusion of roughly half the dataset into Constellations, independent of the lack of strict thresholds for haplotype definition, was the uncurated inclusion of elements identified via the ‘extend’ command within the high confidence set (16). Given current insights into *Starship* biology, we hypothesize that these "extended" elements may represent or be associated with SLRs (9,17). However, validating this hypothesis remains beyond the scope of the present study and warrants dedicated future research. Consequently, we strongly recommend that researchers avoid utilizing the starfish extend command without implementing rigorous downstream filters. Alternatively, researchers should employ stargraph (17) to gain a more comprehensive understanding of regions associated with putative *Starship*-associated tyrosine recombinases that escape confident detection by the starfish “Element Finder” module (4).

### 3.5. Overrepresented orthogroups in *Aspergillus fumigatus*

To identify common genes within the complete set of non-degraded elements in *A. fumigatus*, we performed an overrepresentation analysis on a subset of sequences containing less than 5% degenerated bases (Ns). This analysis identified 36 orthogroups represented across approximately 25% to 41% of the elements. To filter out orthogroups whose overrepresentation was driven primarily by a single *navis*, we performed an enrichment analysis focused on *Tardis* and *Osiris*, which represent the most heavily populated *naves* in the dataset. This revealed that 11 of the 36 orthogroups were significantly enriched in *Osiris*, while 12 were enriched in *Tardis*, leaving only 13 overrepresented orthogroups that were more broadly distributed across diverse *navis* lineages. Eight of these 13 overrepresented orthogroups were functionally annotated as FRE, *Spok* homologs (a toxin-antidote protein), DUF3723, protein kinase-like proteins, PLP, NLR, Fido domain-containing protein, and aminoglycoside phosphotransferase. Among these, five correspond to previously established auxiliary genes (7). Meanwhile, the remaining three orthogroups with functional annotations correspond to two phosphotransferases and one putative protein from the Fic/Doc family (81).

Interestingly, the putative protein from the Fic/Doc family (AFUA_8G06180 locus in *A. fumigatus* Af293) has been reported to increase its expression during conidial formation (82,83). The precise evolutionary and functional relevance of these genes to *Starship* biology remains an open question that might be of interest in future research.

### 3.6. *Starship* synteny is strongly related to taxonomy in the starbase dataset

After validating the pipeline performance, we expanded our analysis to the starbase v0.3.0-pre dataset (71), which comprises a diverse collection of 2071 dereplicated elements from various classes within Pezizomycotina, to demonstrate the pipeline utility. Using host taxonomy availability as our primary inclusion criterion, we first evaluated the network topology based on the host genus. The resulting network assortativity ranged from 0.8845 to 0.991, indicating a strong preference for connections between elements within the same genus (Figure 5; Supplementary data).

**Figure 5.**
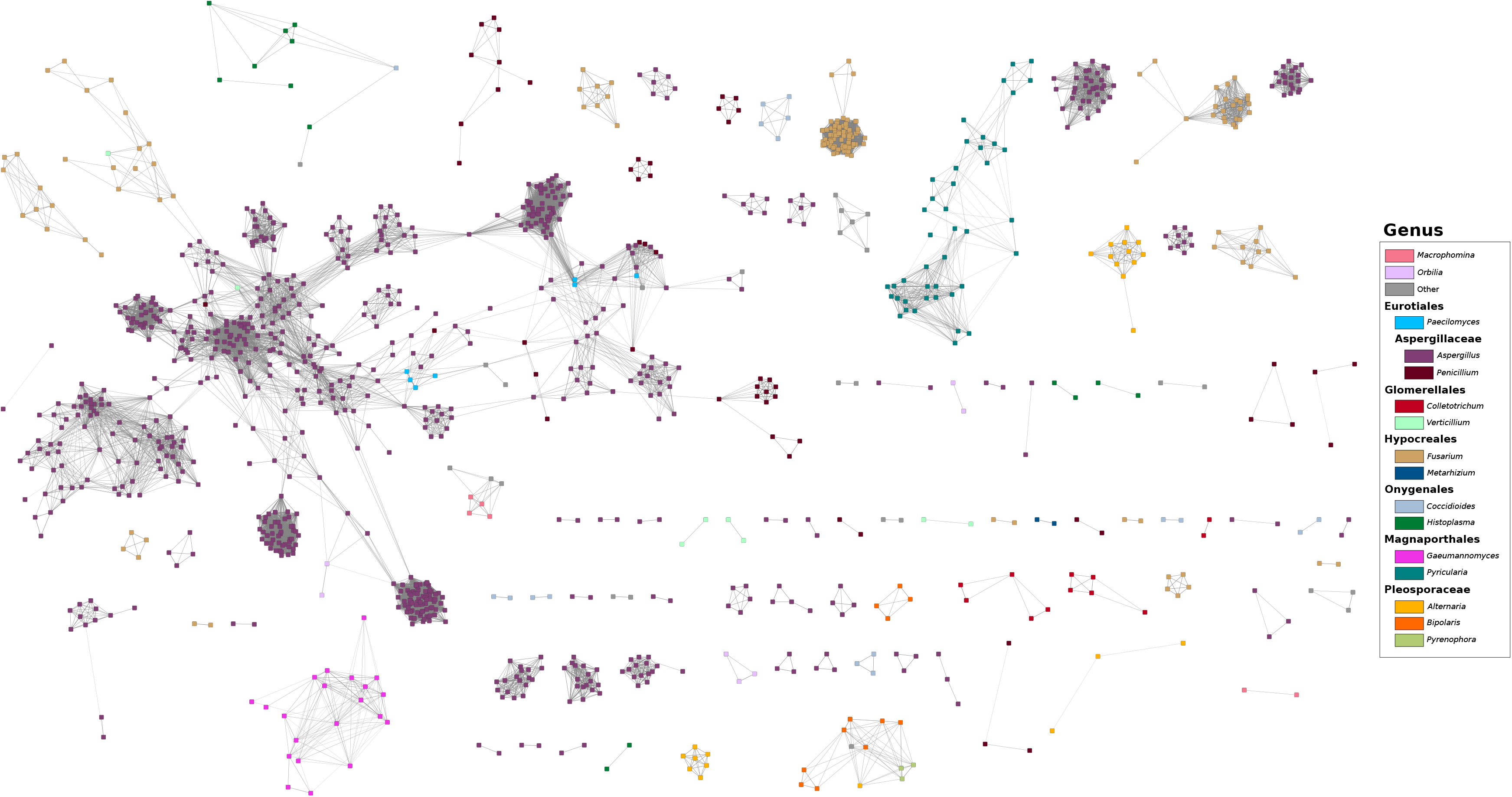
**StarCrew cargo dynamic analysis using the starbase dataset**. Results shown are derived from the preprocessing subset with a minimum gene content of 5. Graph representations generated using a Prefuse force-directed layout illustrating syntenic-based clustering using five anchor points, e-value of 1e^-2^ with metric-based filtering strategy. Node color represents the Genus of the host genome. Genus legend is organized by shared Family and Order.

Inter-genus connections were primarily confined to elements within the same taxonomic family, with a small subset of these relationships persisting even under stringent parameters (e.g. shared elements between *Aspergillus* and *Penicillium*, or *Bipolaris* and *Exserohilum*). Conversely, connections between elements belonging to different orders (and rarely, different classes) were consistently disrupted as stringency increased. This suggests that most connections beyond the taxonomy family level are driven by a limited set of shared genes rather than broad syntenic conservation. These findings align with previous reports where *Starship* mobilization appears to be phylogenetically constrained (11,35), and some of these cases have been previously identified (5,6).

However, it is worth noting that shared elements between species of the same order have been previously reported and recovered in our network (Figure 5), such as those between *Paecilomyces* and *Aspergillus* (8). Furthermore longer distance transfers have been documented, including shared elements between *Spiromastigoides* and *Aspergillus* (Eurotiales order) (8), which was identified in our network (Figure 5), and shared elements between *Fusarium* and *Verticillium* (Hypocreomycetidae subclass) (14), which were not detected in our network due to the absence of those elements within the Starbase database version utilized in this study.

Since the initial dataset was dereplicated using a type element selection for each SSA (a group ID representing highly similar individual elements), we cross-referenced all elements within each SSA with available host species information (71). Most SSAs were restricted to a single genus, reinforcing our previous findings regarding phylogenetic constraints. The only exception was SSA003616, which was predominantly associated with the genus *Penicillium* but also contained a single documented copy in *Amorphotheca resinae*. Manual inspection was performed to confirm the presence of this shared element across distinct taxonomic classes (detailed methods provided in the Supplementary Information). Specifically, we identified the contigs harboring the *Starship* copy in *A. resinae* strain KUC3009 and *Penicillium chrysogenum* strain R4101. In this case, the contig from *P. rubens* strain 43M1 (corresponding to the host strain of the type sequence) could not be utilized because it was too short for extended synteny evaluation. Nevertheless, dotplot analysis via the YASS web server (48) confirmed that the two contigs share sequence identity exclusively within the region surrounding the element, indicating that this cross-class match is not an artifact of assembly contamination (Supplementary figure S10). Furthermore, we identified that this shared sequence region is ∼10 kb larger than the putative element in starbase, suggesting that the element size is currently underestimated. A quick search for known protein domains in this region with CD-search (84) confirms the presence of a DUF3435 domain near the element boundary, and the additional 10Kb nucleotides appears to be related to non-LTR TEs. Nonetheless, further analyses will be required to definitively confirm this shared *Starship* and fully resolve its boundaries.

### 3.7. Preliminary Functional Constellations of starbase elements

Similar to the *A. fumigatus* dataset, we defined in this case a preliminary set of Constellations from elements currently available in starbase v0.3.0-pre. Specifically, we analyzed a subset of 912 elements from starbase dataset filtered to prioritize highly probable active elements (captain gene with confidence level 3, >5 total genes, <30% RIP-like signal coverage, and >30% GC content). This resulted in 132 Constellations, based on (sub)cluster definition from StarCrew (Supplementary Figure S11). As many of these elements are still uncurated, we selected a single Constellation of interest and expanded the set of representative elements by searching publicly available NCBI genomes (detailed methods provided in the Supplementary Information).

We selected a Constellation representing an inter-order transfer between species within the class Eurotiomycetes (Supplementary Figure S11). This constellation represents a share *Starship* between *Spiromastigoides sp.* (SSB000313) and *Aspergillus spp.* (SSB003670 and SSB003716), which was previously reported (8). Here, we recover additional uncharacterized elements to compile a comprehensive Constellation, designated Ceur FC1 (For Class Eurotiomycetes). This search identified a Constellation comprising 13 non-degraded elements distributed across multiple species of *Aspergillus* and a single ocurrence in *Spiromastigoides* (Supplementary table S5, Supplementary Figure S12). Furthermore, during our search across publicly available NCBI genomes, we identified a structural rearrangement within an element in *A. versicolor* strain D5, which was subsequently incorporated as a degraded copy (Supplementary Table S5). Additionally, the element in *A. versicolor* ATCC 9577 was incorporated later and is currently flagged as degraded due to a scaffold break that does not allow us to accurate identified the *Starship* terminal boundary (Supplementary Table S5). The corresponding nucleotide FASTA sequence file is provided in the Supplementary Information. The Constellation was automatically partitioned into two STARs; however, owing to sequence uncertainty within the assembly gap between joined contigs across a subset of elements, this group was designated as a third STAR manually (Supplementary Table S5). Regarding core gene composition, 34 orthogroups were identified as core, including the captain gene, auxiliary genes, and protein domains associated with other TEs, among others. Within this Constellation, two core genes of interest potentially conferring beneficial traits include a putative sodium/calcium exchanger protein and a putative copper-translocating P-type ATPase.

### 3.8. Nesting identification is sensitive to element boundaries

Beyond clustering and core gene identification, the pipeline is capable of identifying nesting events where one element is integrated within another (Figure 6A,B). While a similar feature is available in starfish (4), StarCrew offers the distinct advantage of allowing users to cross-analyze elements aggregated across multiple independent starfish runs. However, we observed false-positive nesting events in instances where element boundary definitions were probably imprecise (Figure 6C, D). While this was not the primary intended purpose of the nesting function, these occurrences offer significant diagnostic value for researchers during data curation. Specifically, these false positives highlight discrepancies in boundary identification across different copies of the same element, serving as a flag for manual refinement. This unexpected utility can facilitate a more accurate identification of direct repeats (DRs) and the characterization of target sites.

**Figure 6.**
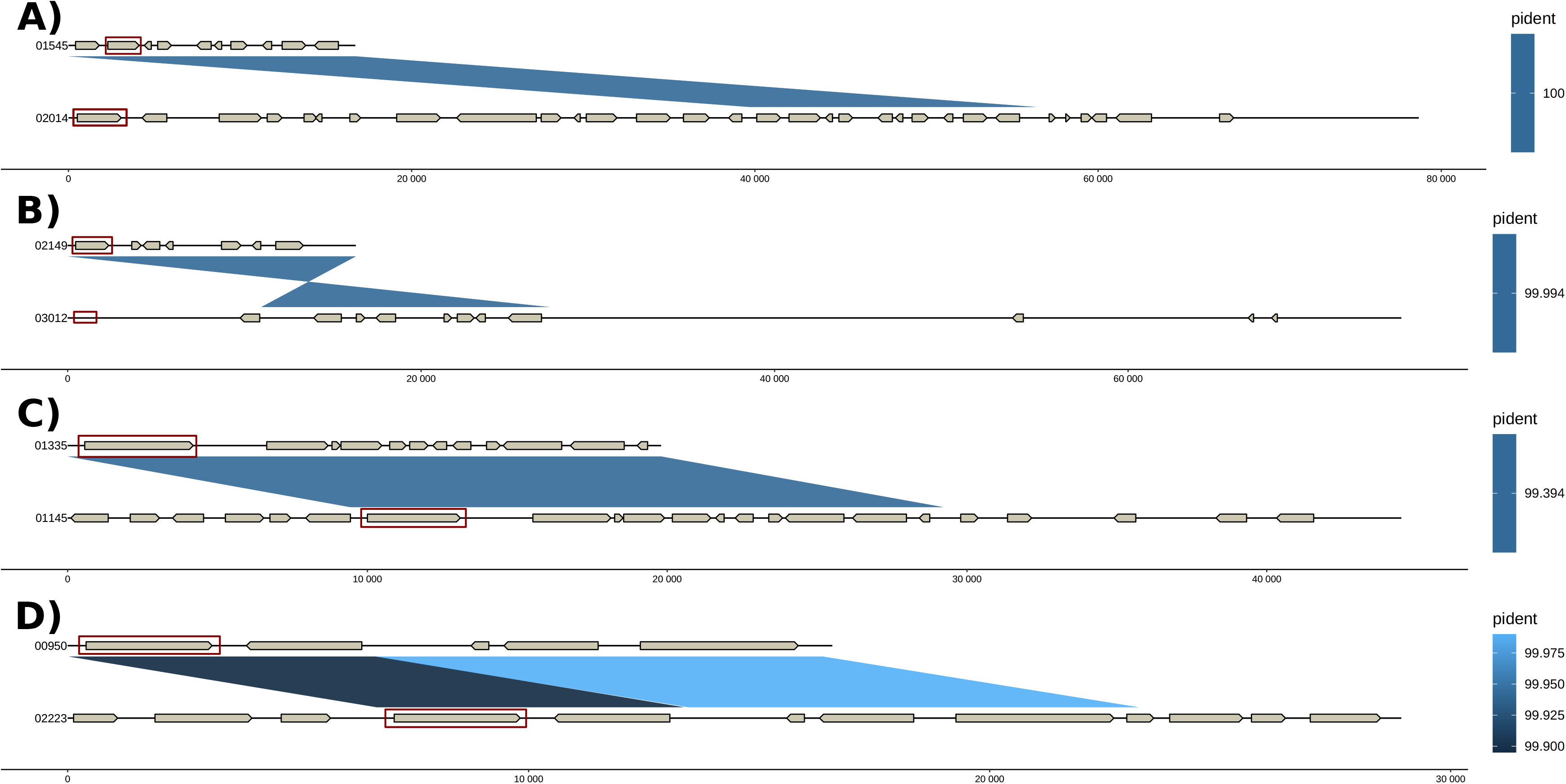
Representation of nesting events identified by StarCrew in the starbase dataset. Nucleotide synteny visualization of putative nesting events. **A, B)** True positive nesting event. **C, D)** False positive nesting events. Red rectangles highlight the position of the identified captain gene/pseudogene within each element by StarCrew. Pident indicates the percentage of nucleotide identity. Mapping of Starbase IDs to StarCrew- generated codes: 01545 (SSB001545); 02014 (SSB002014); 02149 (SSB002149); 01335 (SSB001335); 01145 (SSB001145); 00950 (SSB000950); 02223 (SSB002223).

### 3.9. Filters remove sparse elements rather than specific clades

A potential concern for researchers utilizing this pipeline is whether the application of filters might disproportionately eliminate specific clades of *Starships*, thereby inadvertently reducing the diversity of the dataset. To evaluate this, we constructed a maximum-likelihood phylogeny of captain protein sequences across three distinct filter configurations (Figure 7). The resulting topology demonstrates that excluded elements are broadly distributed across the entire phylogenetic landscape rather than being concentrated within a specific lineage. Additionally, topological discordances in *Starships* family relations with the previously established captain phylogeny (4) can be attributed to both the presence of pseudogenes and gene fragments in our initial dataset and differences in tree inference methods.

**Figure 7.**
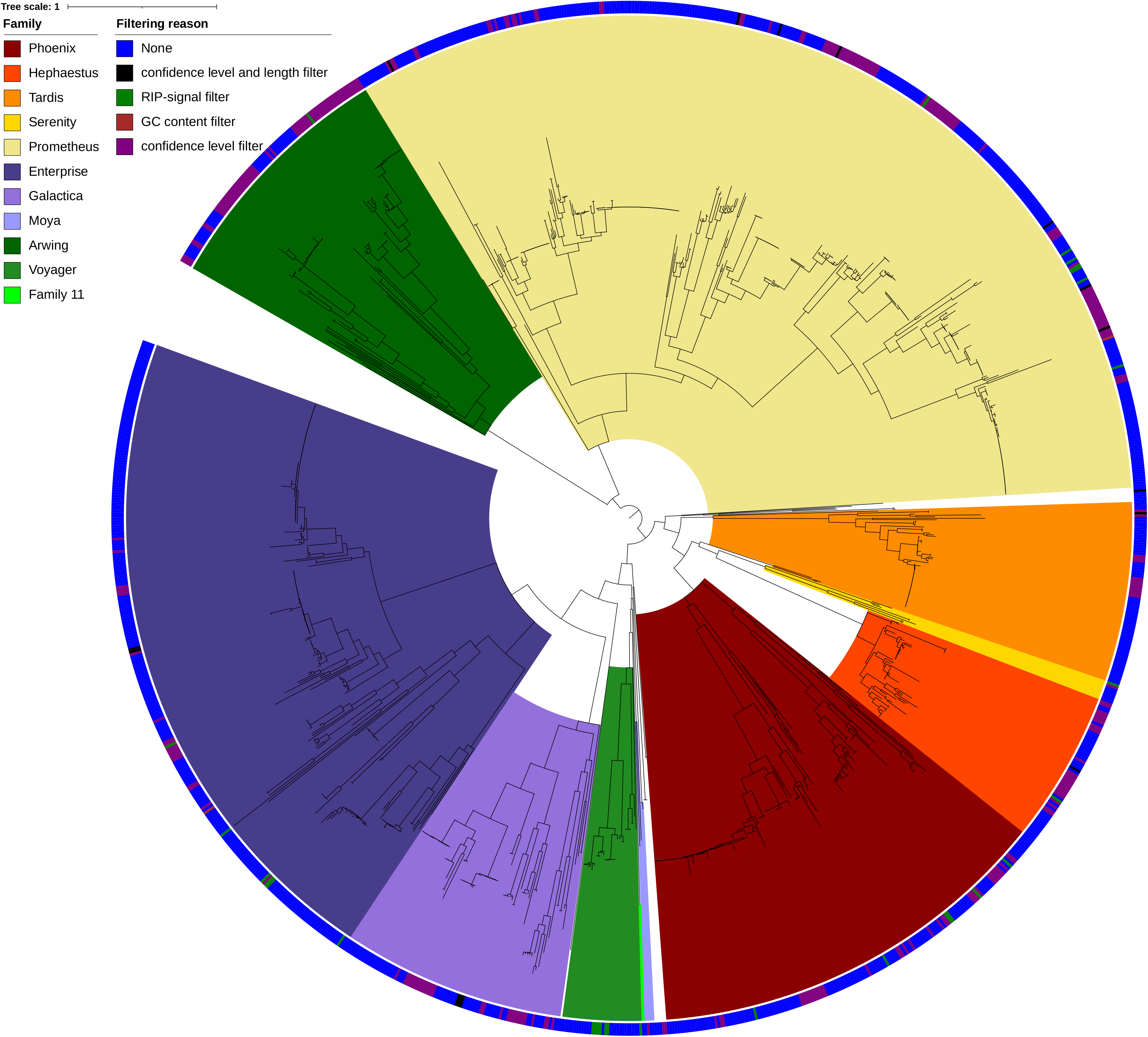
Phylogenetic tree of captain proteins from dereplicated starbase elements. The tree was inferred using Maximum-likelihood analysis with 1,000 SH-aLRT and 1,000 ultrafast bootstrap (UFboot) iterations. All near-zero length branches have been collapsed. The tree is rooted in a position that allows comparison with previously stablished Starships captain tree (4). Clades are color coded according to *Starship* family. The color strips at the tips indicate the specific filtering criteria for which an element was flagged; ‘None’ denotes elements that passed all tested parameters.

While some instances appeared to suggest the removal of entire clades (Figure 7), a closer inspection revealed that these events were driven by data quality rather than phylogenetic bias. By examining five excluded clades comprising more than 10 elements each, we observed that these consisted of redundant elements with structural variations (Supplementary Figure S13). Furthermore, all such elements originated from the uncurated portion of the dataset (71); this is consistent with the filters identifying and excluding potentially inactive or degraded elements, rather than a broad loss of diversity. This trend may be particularly useful for users focused on identifying the most probable active *Starships*. Interestingly, when evaluating the elements that failed to retain a suitable captain gene under the most stringent thresholds, the pipeline was also unable to recover a corresponding pseudogene for the majority of these candidates. This reinforces our benchmarking observation that pseudogene identification may lack the sensitivity to recover fragmented sequences. However, to the best of our knowledge, there’s no alternative pseudogene detection approach that does not rely on direct nucleotide identity to known genes.

Critically, the filtering process does not appear to target specific captain families; representatives of all initially described families (4) remain present even under the highest stringency (Figure 7). In summary, the sparse distribution of excluded elements across the tree supports the conclusion that StarCrew filters do not compromise the overall phylogenetic range of *Starship* diversity.

### 3.10. Auxiliary genes successfully identified as overrepresented in *Starships*

To evaluate the utility of the orthogroup overrepresentation module across diverse taxa, we analyzed a subset of 912 elements from starbase dataset filtered to prioritize highly probable active elements as described also for the Constellation definition. In this dataset, we identified 42 overrepresented orthogroups with frequencies ranging from 7.02% to 35.42%. However, this dataset exhibited a significant taxonomic bias, with *Aspergillus* (predominantly *A. fumigatus)*, representing 69.52% of the elements. To isolate broader evolutionary signatures, we repeated the analysis excluding all *Aspergillus* representatives. This reduced the outlier set to 8 orthogroups, present in 26.25% to 48.20% of the remaining elements. This shift indicates that ∼80% of the initially identified orthogroups are likely specific to *Aspergillus*-associated *Starship* lineages in the analyzed starbase dataset.

The previously established auxiliary genes (5,7) were consistently recovered as outliers in both analyses of the starbase dataset. NLRs were represented across three overrepresented orthogroups in the full dataset (39.47% of elements) versus a single overrepresented orthogroup in the non-*Aspergillus* dataset (33.81% of elements). The remaining auxiliary genes were each identified within single overrepresented orthogroups, including PLP (32.79% full vs. 42.81% non-*Aspergillus*), FRE (33.99% vs. 43.89%), and DUF3723 (30.92% vs. 48.20%). Interestingly, the overall proportion of elements carrying these auxiliary genes was lower than previously reported values (5). We propose two primary factors for this discrepancy: 1) high sequence divergence may distribute these auxiliary genes across multiple distinct orthogroups, similar to patterns observed for the captain gene; and 2) the inclusion of uncurated elements may introduce false positives or non-mobile sequences that artificially dilute the observed frequency of auxiliary genes. In the case of NLRs, the first factor is particularly expected, given that they comprise a highly diverse multidomain group of receptors that are not homologous (85). To test this hypothesis for single-domain auxiliary genes, we evaluated DUF3723 and PLP. We found that DUF3723 domain-containing proteins are distributed across 21 orthogroups in the full dataset (44.62% of elements) and 16 orthogroups in the non- *Aspergillus* dataset (56.47% of elements), whereas PLP domain-containing proteins are distributed across 6 orthogroups in both datasets (33.22% full vs. 43.52% non- *Aspergillus*). These results demonstrate that while outlier analysis highlights the single most frequent orthogroup, auxiliary genes are often dispersed across multiple orthogroups. Nevertheless, even when aggregated, these proportions do not fully reach the previously reported prevalence of >60% for these genes (5). Still, these proteins maintain a high relative prevalence compared to other cargo genes within *Starships*, and the potential inclusion of uncurated, non-mobile, or false-positive elements likely accounts for this remaining deficit.

Notably, Myb/SANT domain-containing proteins were not highlighted as overrepresented in this broad analysis, despite prior reports of their prevalence in the 3’ regions of elements (9). Other significantly overrepresented orthogroups included *Spok* homologs, a putative protein kinase, and protein domains associated with other TEs (supplementary data).

### 3.11. A putative pseudokinase as a conserved gene in *Starships*

In addition to the previously described auxiliary genes, a notable overrepresented gene was a specific putative protein kinase orthogroup, which exhibited high prevalence across both the full (35.41%) and non-*Aspergillus* (35.61%) datasets. This orthogroup is distributed broadly across the captain phylogeny (Figure 8), suggesting that it may represent a conserved component across a wide range of *Starship* elements. Indeed, with the only exception of *Arwing* and Family 11, at least one representative from every *Starship* family contains a gene belonging to this orthogroup (Figure 8). Notably, this specific orthogroup was also overrepresented in the *A. fumigatus* benchmark dataset. In the Af293 reference strain, this orthogroup corresponds to locus tags AFUA_1G16060 and AFUA_5G14970, which are harbored within two elements belonging to the *Nebuchadnezzar navis*. Querying these homologs against the STRING database v12.0 (86) revealed two intriguing patterns: 1) homologs of these genes appear to be restricted to the subphylum Pezizomycotina, and 2) these homologs co-occur with homologs of FRE and DUF3723-domain containing protein (two previously established auxiliary genes (5,7)). Additionally, the locus AFUA_1G16060 has been reported to be regulated by the regulator of G-protein signaling (RgsC) protein that is associated with inorganic and simple nitrogen sensing and/or transport (87).

**Figure 8.**
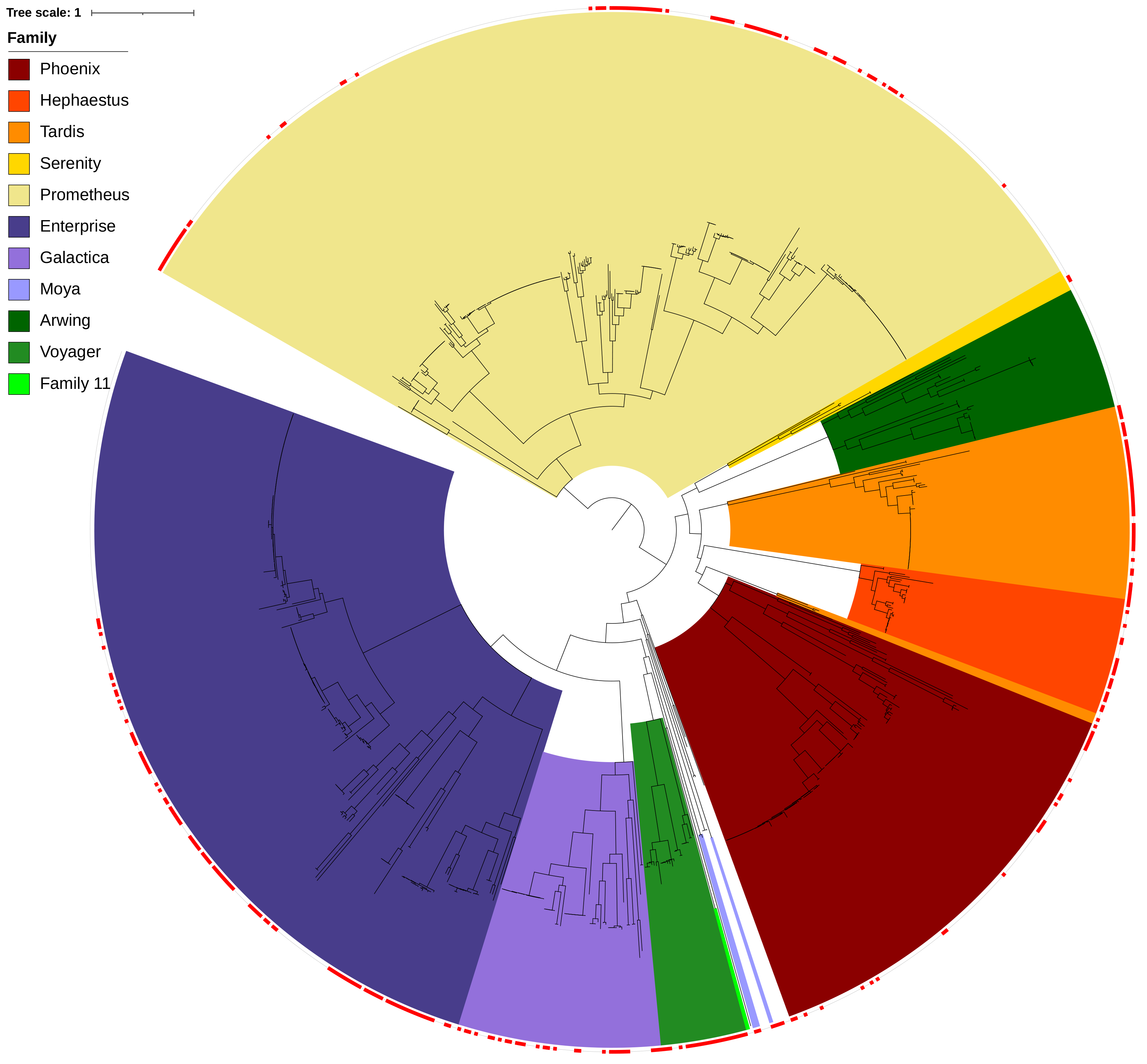
Distribution of overrepresented protein kinase orthogroup among elements with high-confidence captains. The phylogenetic tree was inferred using Maximum-likelihood analysis with 1,000 SH-aLRT and 1,000 ultrafast bootstrap (UFboot) iterations. All near-zero length branches have been collapsed. The tree is rooted in a position that allows comparison with previously stablished Starships captain tree (4). Clades are color coded according to *Starship* family. Red squares indicate the presence of the overrepresented protein kinase orthogroup within specific elements.

We evaluated existing gene models using RNA-seq data from BioProject PRJNA658306 (Supplementary information), which revealed that current annotations did not match the transcriptomic evidence. We update both gene models to align with the RNA-seq data (Supplementary Figure S14), resulting in updated protein sequences. Notably, because the transcriptomic data indicated potential alternative splicing events in both copies, introns were selected based on the abundance of uniquely mapped junction-spanning reads and the preservation of a shared intron–exon architecture across both copies (Supplementary Figure S14). InterProScan (64) analysis confirmed that these revised sequences retain the protein kinase domain. To determine whether this protein corresponds to an active protein kinase or a pseudokinase, we evaluated two lines of evidence: structural topology and sequence motif conservation. Structurally, the protein retains 12 of the 13 conserve subdomains (88,89), lacking only subdomain VIII (which contains the APE motif). At the sequence level, the glycine-rich loop, VAIK motif, catalytic loop HRD motif, and activation segment DFG and APE motifs (89) are absent. Furthermore, structural alignment of the AlphaFold-predicted (90) model for the updated AFUA_1G16060 protein against the *Aspergillus nidulans* calcium/calmodulin-dependent protein kinase cmkA (AlphaFold DB ID: AF-Q00771-F1) suggest a conserved protein kinase fold despite the absent subdomain VIII (Supplementary Figure S15). Together, this evidence suggests that this orthogroup represents a pseudokinase instead of an active protein kinase.

The prevalence of this protein in *Starship* elements (at levels comparable to previously established auxiliary genes) together with its co-occurrence with other auxiliary genes, suggests that this putative pseudokinase is intimately associated with *Starships*, having been overlooked in previous studies (5,7,9). Consequently, this raises an important question for future research: what is the function of this protein within *Starship* biology? Even when pseudokinases lack catalytic activity, they still play an important role in signaling pathways (89,91); therefore, a compelling avenue for subsequent research will be to explore whether this gene participates in signaling pathways that culminate into the activation of element mobility. Although direct evidence supporting this hypothesis is currently lacking, elucidating the role of this highly conserved putative pseudokinase remains an intriguing open question in *Starship* research.

In summary, the orthogroup overrepresentation analysis serves as a powerful exploratory tool for identifying shared genes within a dataset, as demonstrated here by our discovery of a previously overlooked conserved gene. However, users should be aware that results remain sensitive to dataset composition and sequence divergence. We therefore recommend utilizing this module as an initial screening tool for candidate gene discovery, followed by downstream validation using complementary bioinformatic workflows as it was performed for the conserve putative pseudokinase.

## 4. Conclusions

In this work, we present StarCrew, a versatile pipeline designed for the systematic analysis of *Starship* cargo, alongside a new system to group elements with potentially shared beneficial traits. Through our tests, we demonstrate that StarCrew robustly handles datasets with varying levels of heterogeneity, allowing users to fine-tune filtering parameters without compromising the diversity of their data. The pipeline facilitates a broad range of downstream analyses and aids in identifying elements that require boundary curation, providing a valuable diagnostic layer for improving dataset quality. However, the efficacy of the pipeline remains inherently sensitive to the quality of the input data, primarily suboptimal gene models and imprecise boundary delineations. As such, the capabilities of StarCrew are best utilized as part of an iterative analytical process in tandem with the manual curation of low-quality elements. StarCrew uncovered a highly conserved putative pseudokinase across diverse *Starships*, presenting a compelling target for future functional studies. Furthermore, we demonstrate the practical utility of our complementary grouping system by achieving a well-defined set of Constellations of the *A. fumigatus* dataset.

*Starship* identification and curation is a difficult task due to a combination of the fact that *Starship* sequences themselves resemble the background genomes they reside in, with little to no structural features, and that any given genome often has more captain proteins than intact *Starships* due to historic insertion and degradation dynamics. While the starfish tool provides a vital resource for this work, it is highly prone to false positives that require manual attention to eliminate (4). The stargraph tool allows for the exploration of the influence *Starship* insertions have had on a population, and identification of regions that are likely *Starship*-derived (17) but again is limited in its ability to accurately define active, mobile elements. The StarCrew pipeline provides a powerful, cargo-centric perspective for *Starship* curation, which should facilitate the identification of genes that are mobile within fungal species/populations. As mobility was not known to be an inherent trait of fungal genes prior to the discovery of *Starships* (6,7), identifying and categorizing them is crucial to furthering our understanding of how genome dynamics impact fungal evolution.

Finally, as the study of *Starships* is still in the early stages, much is yet to be discovered regarding the evolutionary dynamics and functional benefits of their diverse cargo repertoires. We anticipate that as the scientific community uncovers new structural and biological patterns, the methodologies and filtering strategies integrated into StarCrew will continue to evolve. Ultimately, we hope StarCrew serves as a practical resource for the continued exploration of these unique fungal mobile elements.

## 5. Availability and requirements

**Project name**: StarCrew.

**Project home page**: https://github.com/andres2901/StarCrew

**Operating system**: Linux.

**Programming language**: Bash, R and Python.

**Other requirements**: Anaconda and git.

**License**: GNU AFFERO v3.

## 6. Data availability

All Supporting data regarding the results of the benchmark tests and case studies performed during this project are stored on Zenodo (https://doi.org/10.5281/zenodo.23417113). Supplementary material, Ceur FC1 nucleotide data, and figures are available in the supplementary information file.

## 7. List of abbreviations

Aa: amino acids.
BGC: Biosynthetic gene cluster.
CDS: Coding DNA Sequence.
CME: Cargo-mobilizing mobile element.
DR: Direct Repeat.
DUF: Domain of Unknown Function.
ECP: Element Collinearity Percentage.
FC: Functional Constellation.
FRE: Ferric reductase.
GCP: General Collinearity Percentage.
HMM: Hidden Markov Model.
NLR: NOD-Like receptor.
IQR: Interquartile range.
MC: Medcouple.
NLR: NOD-like receptor.
LTR: Long terminal repeat
PDB: Protein Data Bank.
PLP: Patatin-like phospholipase.
RIP: Repeat-Induced Point mutation.
SLR: *Starship*-like region
Spok: Spore Killer.
*ssf*: Starship formaldehyde cluster.
SSA: Group accession tag.
SSB: Individual Ship accession tag.
SSP: Strong Syntenic Pair.
TE: Transposable element.
TIR: Terminal Inverted Repeat.
UF: Upper Fence.
UPGMA: Unweighted Pair Group Method with Arithmetic Mean.
YR: tyrosine recombinase.

## 8. Ethics declaration

### 8.1. Ethics approval and consent to participate

Not applicable.

### 8.2. Consent for publication

Not applicable.

## **10.** Author’s contribution

Andres Felipe Lizcano-Salas: Study Conceptualization, pipeline Design and Development, Validation, Formal Analysis, Data Curation, Writing – Original Draft. Martin Ryberg: Writing – Review and Editing. Aaron A. Vogan: Study Conceptualization, Data Curation, Supervision, Writing – Review and Editing. All authors have read and approved the final manuscript.

## Acknowledgment

The authors thank Dr. Anastasios Samaras, Dr. Emile Gluck-Thaler and Dr. Samuel O’Donnell for providing feedback under the early development of the synteny clustering approach, as well as Dr. Adrian Forsythe for providing early access to starbase dataset and related metadata. Finally, we would like to thank Adam Taranto, Dr. Andrew Urquhart, and Dr. Emile Gluck-Thaler for their feedback on the manuscript.

