## Supplementary material for "StarCrew: A pipeline for exploratory analysis of *Starship* cargo": Supplemetary_information_file: Supplementary information.docx

**Supplementary information: “StarCrew: A pipeline for exploratory analysis of *Starship* cargo allowed the identification of a conserved pseudokinase”**

Lizcano-Salas, Andres Felipe; Ryberg, Martin; Vogan, Aaron A.

**Content**

### **Supplementary methods**

#### 1. **Initial test of Clustering based on collinear regions**

This test was initially designed to cluster elements to posteriorly identify putative movement events. Consequently, the analysis was highly focused on identifying conserved chunks of DNA that potentially represent these mobilization events. This objective was the guide for the "true positive" definition established in the "1.2 Synteny analysis test" section; however, that specific definition became outdated when the project's goal shifted toward developing a community-available wrapper.

##### **1.1 Test Dataset**

Two datasets were utilized for this initial test: a ‘curated’ dataset containing 229 elements and an ‘un-curated’ dataset with 607 elements, both including *Starship* family association information. The curated dataset is entirely nested within the larger, un-curated dataset. Initial testing and parameter optimization for synteny analysis were performed exclusively on the curated dataset, which was partitioned into families for individual analysis. The un-curated dataset was subsequently used to assess the impact of including low-confidence data on the results and to identify behaviors that can be only observable in a larger data sample.

To prevent discrepancies between elements caused by differences in gene prediction pipelines, we used Metaeuk v6.a5d39d9 (1) to predict the gene content of all elements. Our protein database was constructed from several sources: OrthoDB v12.1 proteins from Pezizomycotina classes (Dothideomycetes, Eurotiomycetes, Leotiomycetes, and Sordariomycetes) (2), proteins shared within the Starfish toolkit (3) (including "captain" and auxiliary genes), our internal database of captain's sequences, UniProt fungal sequences (downloaded between July 7 and 11, 2025), and UniProt bacterial sequences with an annotation score of at least 3 (downloaded on July 15, 2025) (4). This database was then processed to ensure quality and reduce redundancies. Using SeqKit v2.10.0 (5), duplicate sequences were removed and sequences with unknown amino acids ("X"), as well as those shorter than 50 or longer than 10,000 amino acids. A representative set of sequences was then obtained using CD-HIT v4.8.1 (6) with a 95% identity threshold. Only these representatives were used as the protein database for Metaeuk gene prediction. The Metaeuk predictexon module was run with the following parameters: -s 7.5 --exhaustive-search-filter 1 --filter-msa 1 --chain-alignments 1 --remove-tmp-files 1 --max-seqs 500 --use-all-table-starts 1 --start-sens 7.5. Subsequently, the reduceredundancy and unitesetstofasta modules were used to generate a gff file. Incomplete gene models lacking both start and stop codons were removed using AGAT v1.5.1 (7).

A custom script was then used to determine the number of genes, average gene length, and average intergenic length for each element. Finally, the proteome of each *Starship* was extracted using agat_sp_extract_sequences.pl with the -t exon --merge -p options (7). For this test, only elements with at least ten predicted genes were included.

##### **1.2 Synteny analysis test**

For synteny analysis of the elements, we used the interspecies_synteny() function from the Syntenet v1.10.0 R package (8), based on shared collinear genes. During this project, we do not have a set of pairs with "true" synteny, meaning we are only able to detect false positives and true positives through the various tests. Resulting pairs were visually evaluated using the YASS web server (9) with default parameters. A true positive was defined as a pair with a visible diagonal (complete or fragmented, forward or reverse) in the dot-plot. Additionally, the sum of non-overlapping alignments of at least 5Kb had to represent a minimum of 20% of the length of either element. Conversely, a false positive was defined as any pair that did not meet these criteria.

Initially, we ran Syntenet following its tutorial (<https://almeidasilvaf.github.io/syntenet/articles/syntenet.html>), using BLAST (10) instead of DIAMOND (11). The results were normalized as a percentage of collinear genes for each pair, which allowed for direct comparisons. Based on these preliminary results, we selected the Galactica family for parameter optimization because it contained the highest number of syntenic pairs.

Next, we tested two search software options: BLASTp v2.16.0+ (10) and DIAMOND v2.1.12 (11), maintaining the default parameters for the *interspecies_synteny()* function.

Initially, BLASTp was used to evaluate three parameters: e-value (1e-3 and 1e-5), scoring matrix (BLOSUM62, BLOSUM80, and PAM30), and percentage of query coverage (50% and 80%). We performed twelve tests, covering all possible combinations. The results indicated that all parameters affected the outcome, with query coverage being the most significant. Generally, an increase in query coverage led to a reduction in false positives. Based on these findings, we chose an e-value of 1e-5 and conducted a second test, evaluating the three matrices with a more stringent 90% query coverage. This test demonstrated a reduction in false positives and maintained consistency in pairs with high synteny. The PAM30 matrix yielded the lowest number of false positives. Consequently, we selected the following BLASTp parameters for subsequent tests: an e-value of 1e-5, a PAM30 scoring matrix, and 90% query coverage. Afterward, we evaluated two sensitive modes (default and fast) of DIAMOND, applying the same parameters selected for BLASTp. The fast mode effectively removed some low-synteny pairs while preserving the consistency of those with the highest synteny. Overall, DIAMOND produced fewer false positives, making it the preferred search software for our analysis.

As the second step, we tested two parameters of the MCScanX algorithm (12): anchor points and maximum number of gaps. These tests were performed using DIAMOND with the previously selected parameters. While we initially designed a test with four combinations (values of 5 and 10 for both parameters), preliminary results were suboptimal. Consequently, we performed an additional ten tests, evaluating values for both parameters between 5 and 10 to identify the optimal settings. The best results were obtained with 8 anchor points and a maximum of 8 gaps, in combination with an 8% threshold of shared collinear genes.

Afterward, we applied the selected parameters to all families, both individually and collectively, using the curated and un-curated datasets. This was done to determine if the parameters were universally applicable across all families and to identify any changes in previously established pairs. During this process, we identified some false negatives within certain families that had values exceeding the threshold. These false negatives did not cluster at the bottom of the results table. Despite this, these findings allowed us to define what we termed strong syntenic pairs (SSP). A pair was classified as SSP if it had at least 41% shared synteny. For elements with highly discordant lengths, the definition was a minimum of 45% shared genes in one element and a ratio of at least 1.8 in the percentage of shared genes between the two elements.

#### 2. **Gene prediction for benchmark and Starbase analysis**

We identified that gene prediction for such diverse elements was a challenging task, as Metaeuk results often exhibited a high ratio of false positives and pseudogenes. For this reason, we designed a workflow for this purpose for the rest of the tests. Initially, a Metaeuk 5.34c21f2 (1) prediction was conducted to identify proteins with some level of homology to the putative genes in the elements using the same protein database developed for the initial testing stage and the following parameters: -s 7.5 --orf-start-mode 0 --min-length 50 --min-seq-id 0.3 --cov-mode 1 -c 0.50 --remove-tmp-files 1 --max-seqs 1000 --use-all-table-starts 1 --min-aln-len 50. Based on these results, the proteins that were successfully mapped with Metaeuk were compiled and used as the protein database for Braker3 gene prediction with the following parameters: --softmasking_off --downsampling_lamba=0 --gff3 -–fungus --augustus_args “--genemodel=complete --noInFrameStop=true” (13). Gene models lacking both start and stop codons were subsequently removed using AGAT v1.5.1 (7). As anticipated, the "captain" gene was not consistently identified in all cases. Therefore, we employed an approach similar to the Starfish GeneFinder module (3) for captain identification. Briefly, the Starfish database of captain proteins was used to run Metaeuk 7.bba0d80 (1) using the following parameters: s 7.5 --exhaustive-search-filter 1 --filter-msa 1 --chain-alignments 1 --remove-tmp-files 1 --use-all-table-starts 1 --start-sens 7.5 --orf-start-mode 0 -v 0. The HMM profile from Starfish was then applied to remove false-positive models using hmmsearch with a domain e-value of 1e^-3^. Then, both the Braker3 and captain Starfish-like predictions were merged; in cases of overlapping models, the prediction resulting in the longest gene was retained. To further refine the resulting gene models, we applied two filtering criteria: an intron density threshold of 6 introns per Kb of gene, and a maximum limit of 30 introns per gene. Finally, only the longest isoform per gene is maintained. The scripts used for this gene prediction are stored in the ‘aux/’ directory of the StarCrew repository under “BrakerGenePrediction.sh” and “WholeGenePrediction.sh” names.

#### 3. **Ceur FC1 Constellation expansion and definition**

To construct the Ceur FC1 Constellation, we first defined the boundaries of the three copies available in *starbase* by conducting all-versus-all pairwise contig alignments using the YASS web server (9). Common terminal boundaries across all three copies were then manually identified via multiple sequence alignment. Next, we performed web-based BLAST searches of the full element sequences against the NCBI Whole Genome Shotgun (WGS) database (accessed 8 September 2026) using megablast, discontiguous megablast, and blastn (10). Hits exhibiting >80% query coverage were retrieved and inspected to confirm complete element boundaries. In cases where an element was split across a contig break, both contig ends were manually joined with an 10-N spacer before boundary definition. To this set, three additional *Starship* elements from *Aspergillus* and *Spigomastoides* (14) were added to include related elements harboring shared cargo. Gene prediction was then performed as described in the preceding section. Subsequently, StarCrew analysis was executed using a 30% Repeat-Induced Point (RIP) mutation and GC-content filter for initialization, followed by synteny-based clustering in Constellation mode with an 80% nucleotide coverage threshold. Cluster characterization and orthogroup annotation were conducted to identify core genes of interest within this Constellation. Degraded elements were later added to the final set.

#### 4. **Pseudokinase evaluation**

##### **3.1. *Aspergillus fumigatus* gene model evaluation**

Sequencing reads from Sequence Read Archive (SRA) experiments associated with BioProject PRJNA658306 were downloaded using fasterq-dump (15) and individually mapped against the *A. fumigatus* Af293 reference genome (RefSeq accession GCF_000002655.1) using STAR with default parameters (16). Reads mapping to contigs harboring previously reported *Starship* elements (17) were extracted across all individual SRA runs and pooled. This merged read set was re-mapped against the reference genome using STAR under more stringent alignment parameters (--outFilterType BySJout --outFilterMultimapNmax 5 --outFilterMismatchNmax 5 --outSJfilterReads Unique --outSAMmultNmax 1 --alignIntronMax 1000 --alignMatesGapMax 10000). The resulting RNA-seq alignment was visually inspected using Integrative Genomics Viewer (IGV) (18), and the gene models for AFUA_1G16060 and AFUA_5G14970 were manually updated to conform with the transcriptomic evidence and a conserve exon-intron architecture between both copies.

##### **3.2 Kinase domain evaluation**

Using the updated gene models, translated protein sequences were re-analyzed using InterProScan (19). Three-dimensional structural models for both updated protein sequences were generated using AlphaFold3 (20). Based on these structural predictions, we assessed the presence of canonical protein kinase subdomains and conserved primary-sequence motifs (21,22). To evaluate structural fold conservation, the predicted model for AFUA_1G16060 was aligned against the *Aspergillus nidulans* calcium/calmodulin-dependent protein kinase CmkA (AlphaFold DB ID: AF-Q00771-F1) in UCSF ChimeraX (23). Prior to alignment, default energy minimization was performed within UCSF ChimeraX, followed by structural superposition using the Matchmaker tool with Smith-Waterman algorithm (23).

### **Supplementary Figures Legend**

**Supplementary figure S1. StarCrew Benchmark of cargo dynamic analysis using the resistance gene cluster dataset**. Graph representations of syntenic-based clustering across the benchmark dataset with three anchor points and **A)** e-value of 1e^-2^ and Blastn filtering strategy, **B)** e-value of 1e-2 and metric-based filtering strategy, **C)** e-value of 1e^-2^ and Strong Synteny Pairs filtering strategy, **D)** e-value of 1e^-2^ without filtering, and **E)** e-value of 1e^-5^ with metric-based filtering strategy. Node shapes denote subcluster divisions. Node colors indicate the origin set: green (arsenic resistance), blue (ssf set), and gray (random starbase elements). Synteny characterization analysis of the **F)** arsenic cluster and **G)** formaldehyde cluster, derived from the three anchor points, e-value of 1e^-2^ with metric-baseed filtering test.

**Supplementary figure S2. StarCrew cargo synteny plot within the resistance benchmark dataset.** Visualization of the cluster containing elements harboring the *ssf* formaldehyde resistance gene cluster. The plot represents results using three anchor points, e-value of 1e^-2^ with the Strong Synteny Pair (SSP) filtering strategy.

**Supplementary figure S3. StarCrew Benchmark of cargo dynamic analysis with *Aspergillus fumigatus* dataset**. Graph representations of syntenic-based clustering across the benchmark dataset with **A)** five anchor points an e-value of 1e^-2^ with metric-based filtering strategy, **B)** three anchor points an e-value of 1e^-2^ and Blastn filtering strategy, **C)** three anchor points an e-value of 1e^-2^ and Strong Synteny Pairs filtering strategy, **D)** three anchor points an e-value of 1e^-2^ and metric-based filtering strategy, and **E)** default parameters. Node color denotes previous haplotype classification, while node shape indicates the subcluster classification assigned by StarCrew.

**Supplementary figure S4. StarCrew cargo synteny plot of *Osiris* h3 elements.** Visualization of the cluster containing *Osiris* h3 elements, generated using three anchor points, an e-value of 1e^-2^ with the Strong Synteny Pair (SSP) filtering strategy.

**Supplementary figure S5. StarCrew cargo synteny plot of *Logos* h2 subclusters.** Visualization of elements categorized into distinct sub-clusters using three anchor points, an e-value of 1e^-2^ with the Strong Synteny Pair (SSP) filtering strategy. **A)** Subcluster characterized by truncated gene models, and **B)** subcluster containing non-truncated gene models.

**Supplementary figure S6. Janus navis evaluation based on StarCrew cargo gene dynamic analysis.** Results are derived from the *Janus navis* dataset using three anchor points, an e-value of 1e^-2^ with the metric filtering strategy (except where noted). **A)** Graph representation of *Janus* elements. Node color indicates previous haplotype classification; node shape denotes StarCrew subcluster classification. **B)** Synteny plot of selected *Janus* h1 elements. This panel highlights elements classified into two separate clusters. The rectangle identifies three specific elements from the minor *Janus* h1 cluster marked with a rectangle in panel A. **C)** Synteny plot of the "bridge" subcluster between *Janus* h1 and h2 visualized in panel A. **D)** Synteny Plot from the cluster containing *Janus* h2 of the StarCrew cargo gene dynamic analysis using default parameters, the rectangles mark elements that match the two *Janus* h2 sub-clusters in panel A. **E)** Comparative synteny visualization of selected *Janus* h2 elements. The element enclosed in the square corresponds to the specific node pointer out in panel A.

**Supplementary figure S7. *Lamia h2,* h3 and h4 analysis.** *Lamia navis* evaluation. Graph of *Lamia* h2-h4 obtained by StarCrew cargo gene dynamic analysis using **A)** default parameters; **B)** five anchor points, e-value of 1e^-2^ with metric filtering strategy; and **C)** three anchor points, e-value of 1e^-2^ with strong synteny pairs filtering strategy. Node color indicates previous haplotype classification; node shape denotes StarCrew subcluster classification. **D)** Synteny plot of the h3/h4 subcluster. Visualization of the specific subcluster from panel C containing both *Lamia* h3 and h4 nodes, illustrating shared syntenic regions. **E)** Comparative synteny plot of selected *Lamia* h2 and h4 elements. The sequence marked with an arrow corresponds to the specific node identified by an arrow in panels B and C. Colored boxes indicate the original haplotype classifications for each sequence.

**Supplementary figure S8. Synteny-plot between elements belonging to Aspfum FC5S1 and Aspfum FC11S1.** AF293_s00037 belongs to Aspfum FC11S1 while AF100-12_5_s00786 belongs to Aspfum FC5S1.

**Supplementary figure S9. Synteny plot of Aspfum FC7.** Result from StarCrew ClusterCharacterization with “constellation” flag enabled. The rectangle delineates the genes associated with the fumigermin biosynthetic gene cluster. The arrow points out the element considered as Aspfum FC7S2.

**Supplementary figure S10. Dot-plot between *Amorphotheca resinae* and *Penicillium chrysogenum* contigs harboring a shared *Starships* element.** Dot-plot performed in YASS web-server with e-value threshold of 0.001. JACIXU010000013.1 belongs to *A. resinae* and JAMACR010000024.1 belongs to *P. chrysogenum*.

**Supplementary figure S11. Preliminar StarCrew Constellation network from starbase.** Graph representation generated using a Prefuse force-directed layout illustrating synteny-based Constellation definitions across a subset of *starbase* elements. Node colors represent the host genome genus. The connected component enclosed within the black rectangle highlights the Constellation selected for manual curation and downstream analysis.

**Supplementary figure S12. StarCrew cluster characterization of the Functional Constellation Ceur FC1.** Synteny plot displaying the preliminary set of non-degraded elements comprising Ceur FC1. The arrow indicates the element identified in *Spiromastigoides*.

**Supplementary figure S13. Synteny plot of filtered-out clades.** Visualization of synteny from elements that were excluded during analysis, focusing on clades containing more than 10 elements. Squares delineate the boundaries of individual clades. Green arrows indicate elements filtered out based on the Repeat-Induced Point (RIP) mutation signal.

**Supplementary Figure S14. Integrative Genomics Viewer (IGV) visualization of RNA-seq data at putative pseudokinase loci in *Aspergillus fumigatus* strain Af293.** Read coverage and spliced alignment tracks corresponding to the **A)** AFUA_1G16060 and **B)** AFUA_5G14970 locus tags.

**Supplementary Figure S15. Structural alignment between AFUA_1G16060 and *Aspergillus nidulans* CmkA.** UCSF ChimeraX three-dimensional structural superposition showing the AlphaFold-predicted model of AFUA_1G16060 (orange) aligned against *A. nidulans* CmkA (blue; AlphaFold DB ID: AF-Q00771-F1).
