## Supplementary material for "StarCrew: A pipeline for exploratory analysis of *Starship* cargo": Supplemetary_information_file: FigureS1.pdf

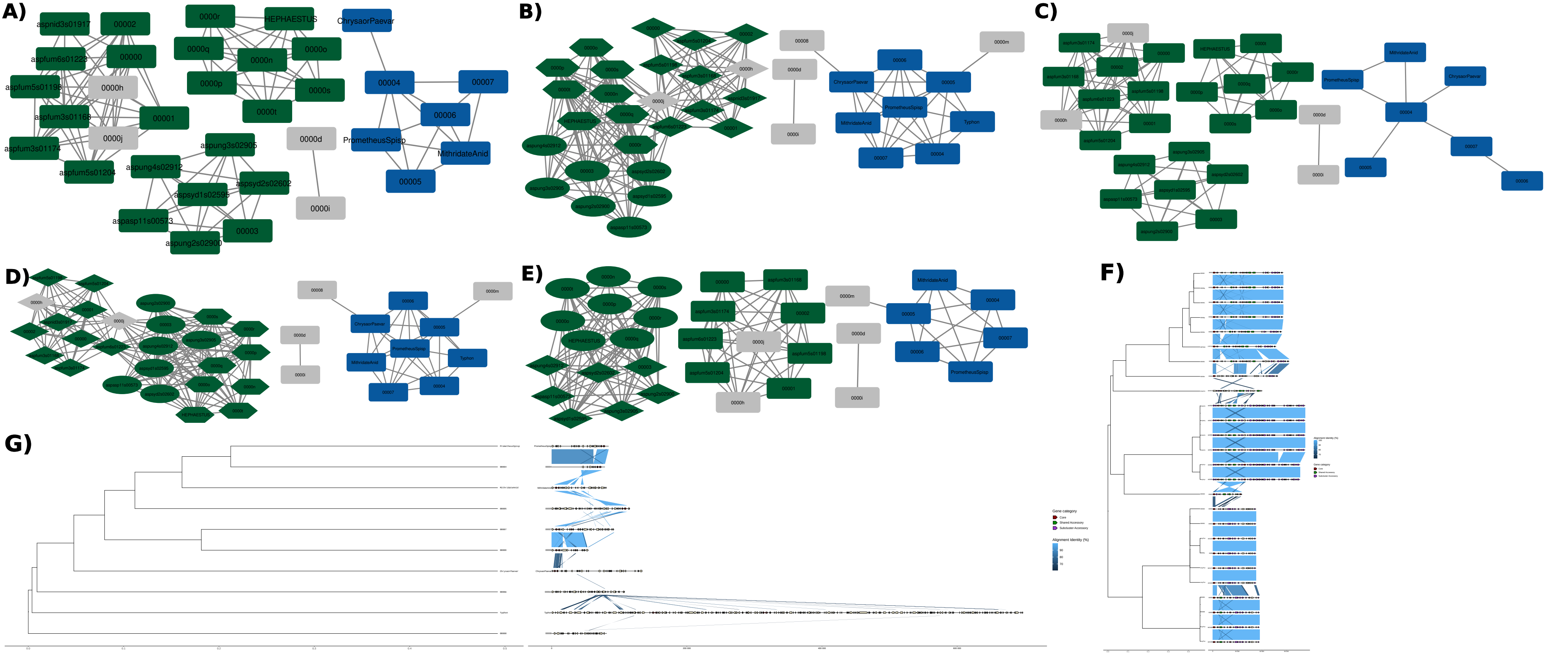
A)
aspnid3s01917
00002
aspfum6s01223
00000
aspfum5s01198
0000h
aspfum3s01168
00001
0000j
aspfum3s01174
aspfum5s01204
0000r
HEPHAESTUS
0000q
0000n
0000o
0000p
0000s
0000t
aspung3s02905
aspung4s02912
aspsyd1s02595
aspsyd2s02602
aspasp11s00573
00003
aspung2s02900
ChrysaorPaevar
00004
00007
00006
PrometheusSpisp
MithridateAnid
00005
0000d
0000i
B)
C)
D)
E)
F)
G)
Gene category
Core
Shared Accessory
Subcluster Accessory
Alignment Identity (%)
