## Supplementary material for "StarCrew: A pipeline for exploratory analysis of *Starship* cargo": Supplemetary_information_file: FigureS2.pdf

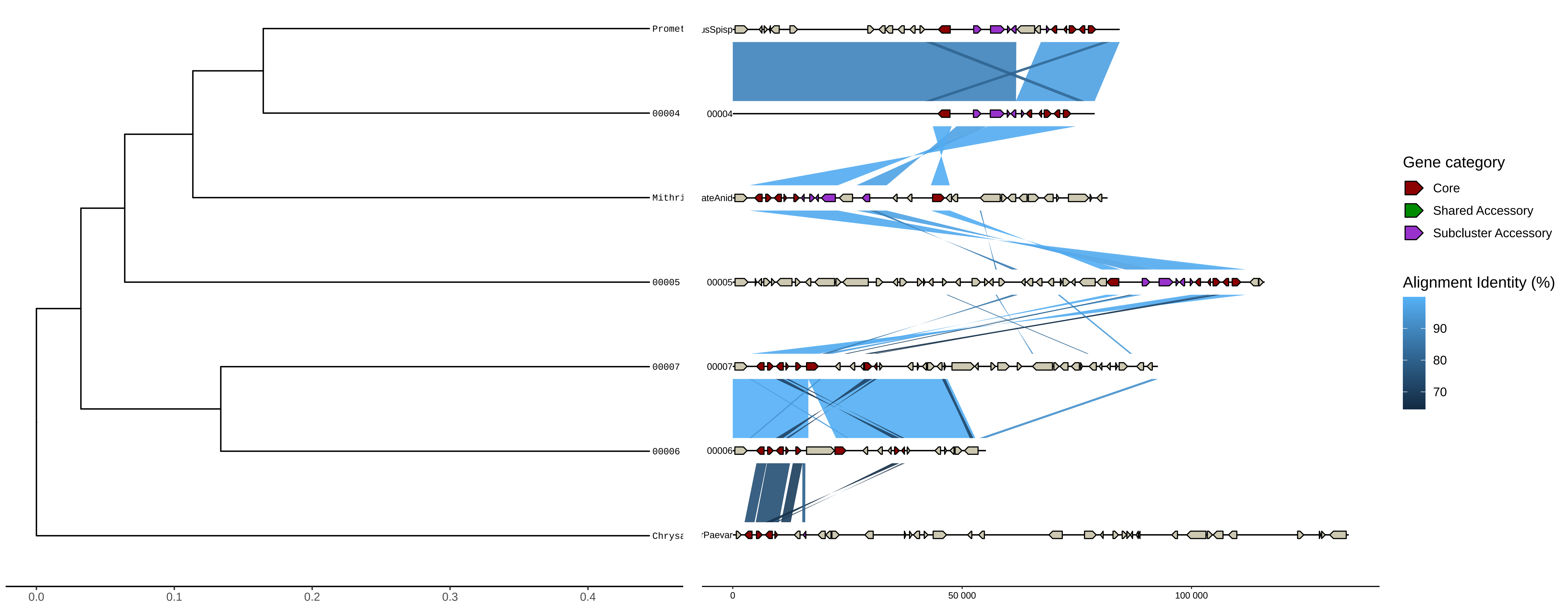

Promet
00004
Mithri
00005
00007
00006
Chrysa
0.0
0.1
0.2
0.3
0.4
usSpisp
00004
ateAnid
00005
00007
00006
rPaevar
0
50 000
100 000
Gene category
Core
Shared Accessory
Subcluster Accessory
Alignment Identity (%)
90
80
70
