## Supplementary material for "StarCrew: A pipeline for exploratory analysis of *Starship* cargo": Supplemetary_information_file: FigureS3.pdf

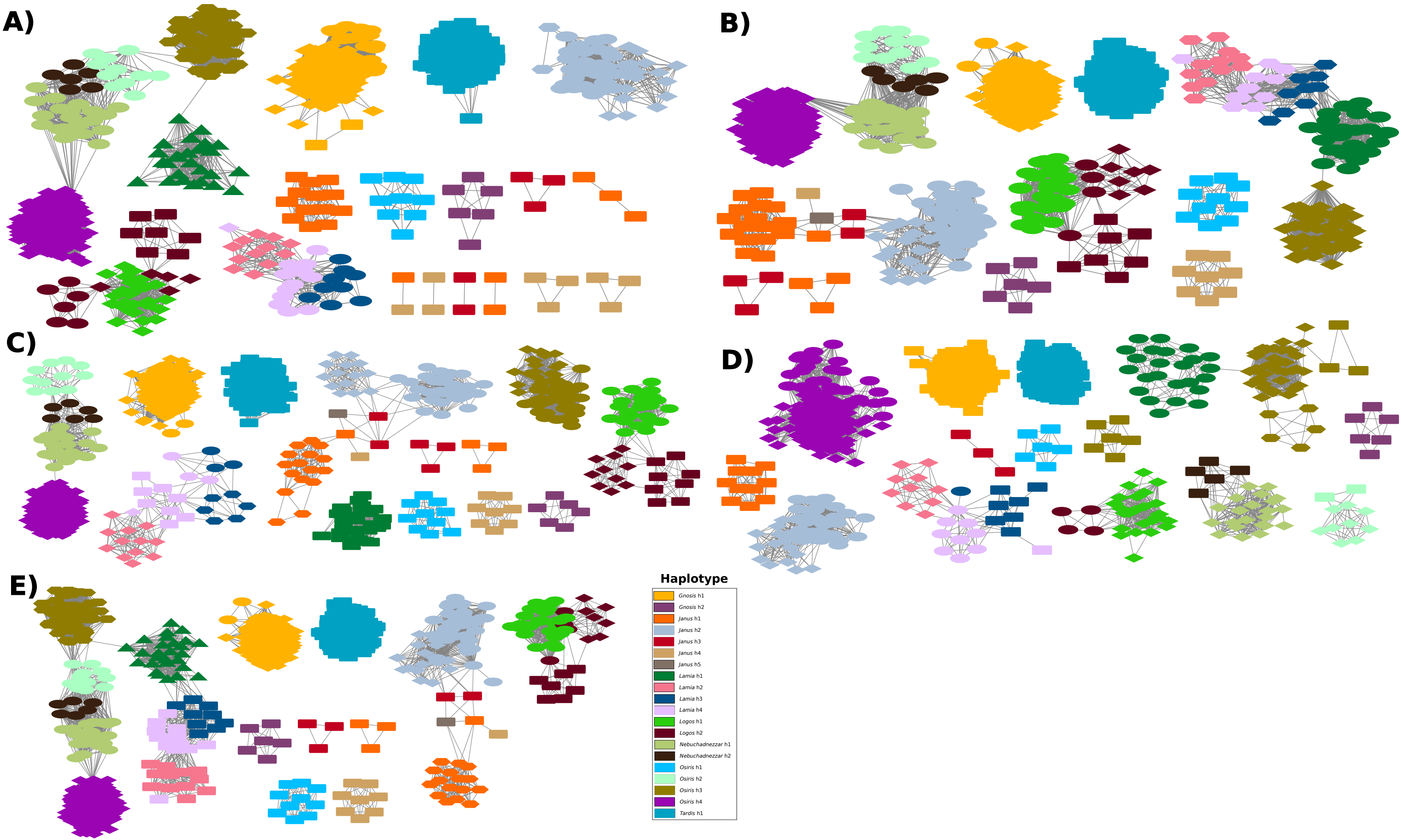
A)
B)
C)
D)
E)
Haplotype
Gnosis h1
Gnosis h2
Janus h1
Janus h2
Janus h3
Janus h4
Janus h5
Lamia h1
Lamia h2
Lamia h3
Lamia h4
Logos h1
Logos h2
Nebuchadnezzar h1
Nebuchadnezzar h2
Osiris h1
Osiris h2
Osiris h3
Osiris h4
Tardis h1
