## Supplementary material for "StarCrew: A pipeline for exploratory analysis of *Starship* cargo": Supplemetary_information_file: FigureS4.pdf

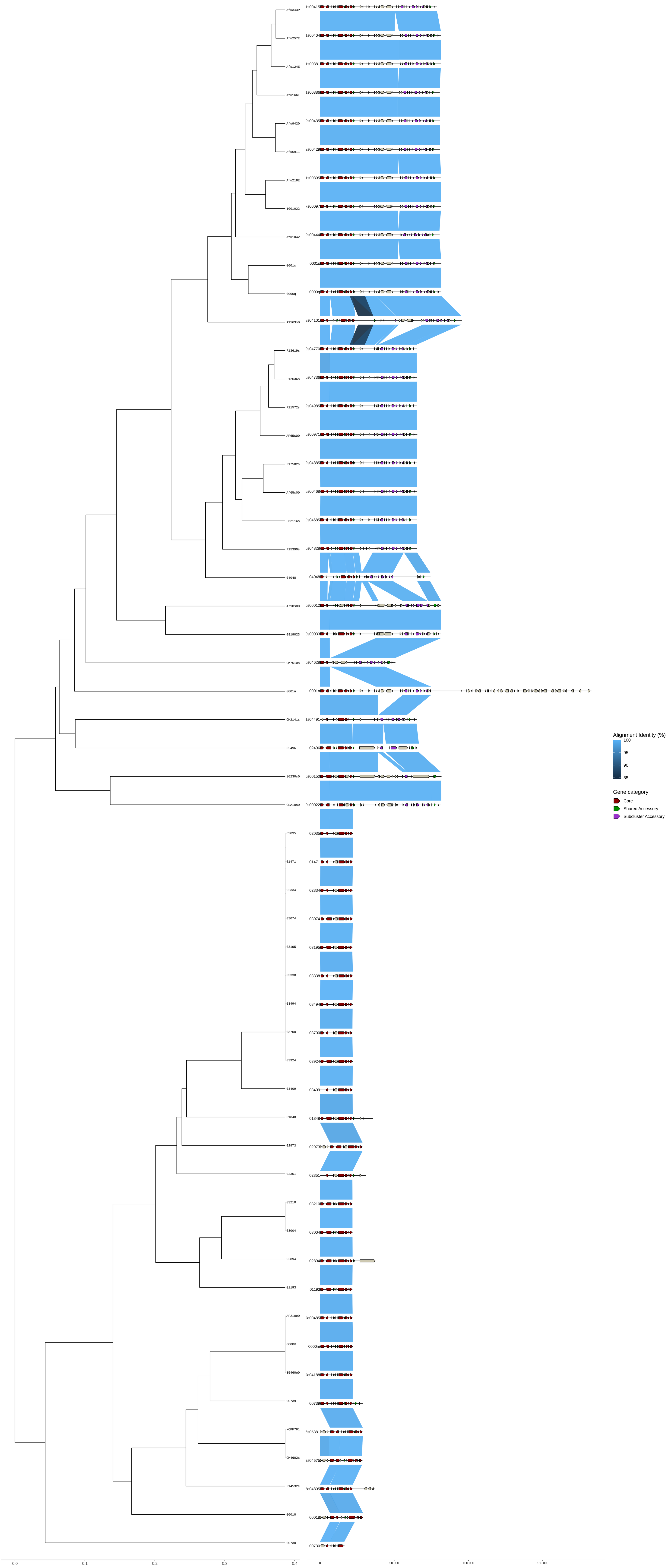

Afu343P
Afu257E
Afu124E
Afu166E
Afu9420
Afu5911
Afu218E
1001022
Afu1042
0001s
0000q
A1163s0
F13619s
F12636s
F21572s
AP65s00
F17582s
Af65s00
F5211Gs
F15390s
04048
4710s00
0819023
CM7510s
0001n
CM2141s
02496
S0230s0
CEA10s0
02035
01471
02334
03074
03195
03338
03494
03700
03924
03409
01848
02973
02351
03210
03004
02894
01193
AF210e0
0000m
B5460e0
00739
NCPF781
CM4602s
F14532e
00018
00730
Ls00415
Ls00404
Ls00381
Ls00386
9s00435
2s00429
Ls00395
7s00097
9s00444
0001s
0000q
3s04101
9s04770
6s04736
2s04985
5s00971
2s04885
5s00468
6s04685
0s04828
04048
0s00012
0s00033
0s04628
0001n
Ls04491
02496
0s00150
0s00022
0.0
0.1
0.2
0.3
0.4
0
50 000
100 000
150 000
Alignment Identity (%)
100
95
90
85
Gene category
Core
Shared Accessory
Subcluster Accessory
