## Supplementary material for "StarCrew: A pipeline for exploratory analysis of *Starship* cargo": Supplemetary_information_file: FigureS5.pdf

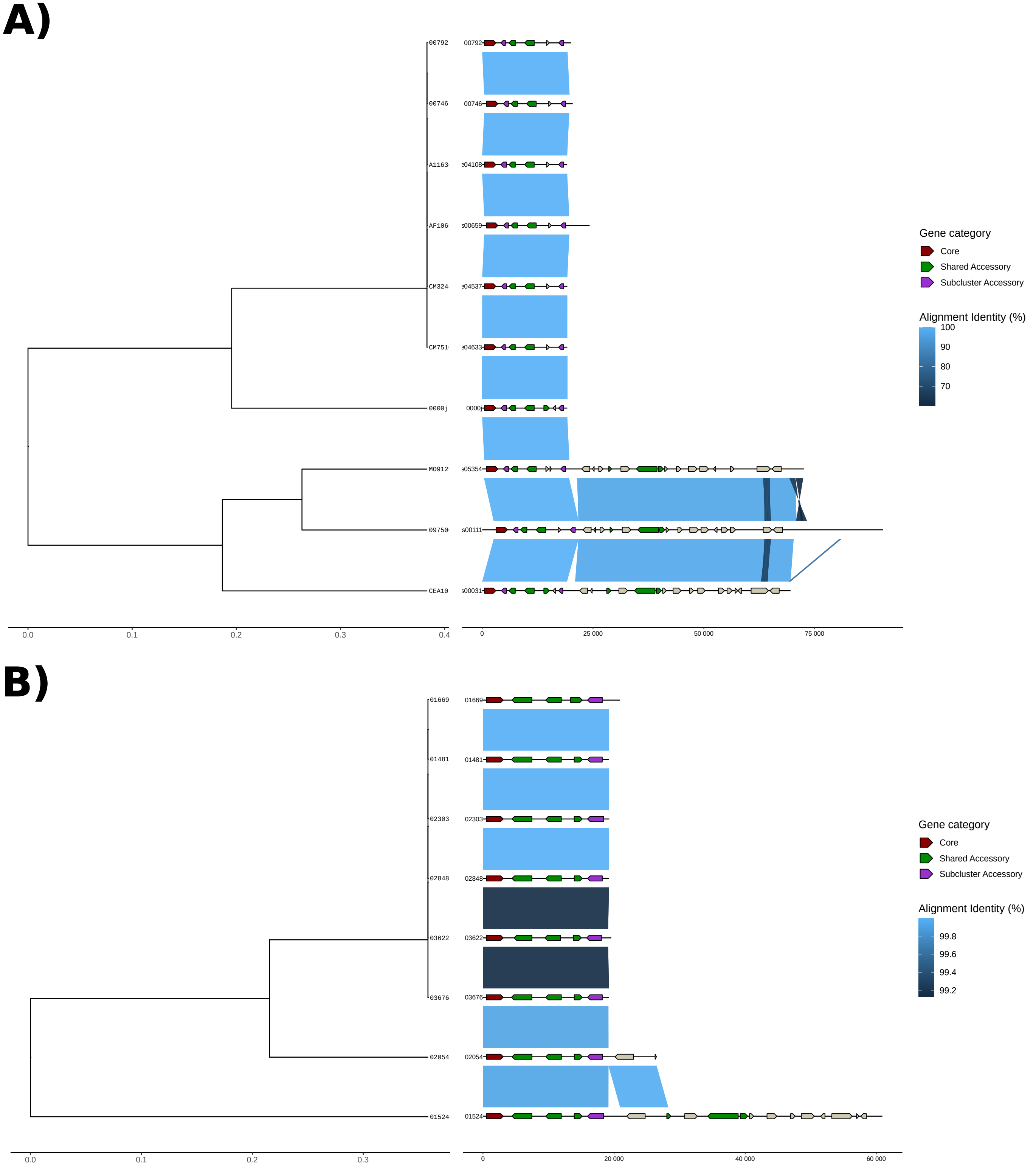
A)
00792
00746
A1163
AF106
CM324
CM751
0000j
M0912
09750
CEA10
00792
00746
e04108
s00659
e04537
e04633
0000j
s05354
s00111
s00031
0.0
0.1
0.2
0.3
0.4
0
25 000
50 000
75 000
Gene category
Core
Shared Accessory
Subcluster Accessory
Alignment Identity (%)
100
90
80
70
B)
01669
01481
02303
02848
03622
03676
02054
01524
01669
01481
02303
02848
03622
03676
02054
01524
0.0
0.1
0.2
0.3
0
20 000
40 000
60 000
Gene category
Core
Shared Accessory
Subcluster Accessory
Alignment Identity (%)
99.8
99.6
99.4
99.2
