## Supplementary material for "StarCrew: A pipeline for exploratory analysis of *Starship* cargo": Supplemetary_information_file: FigureS6.pdf

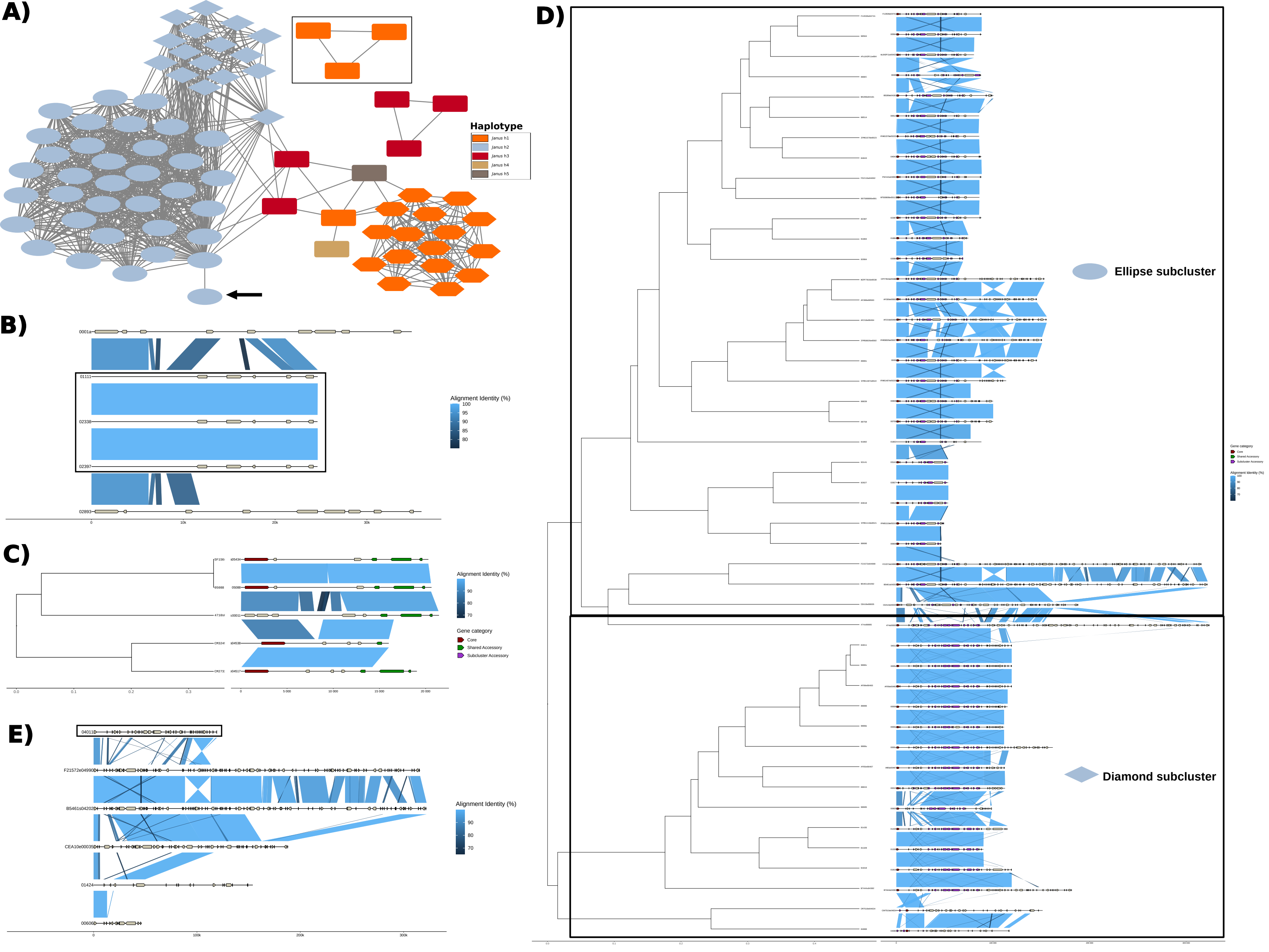
A)
Haplotype
Janus h1
Janus h2
Janus h3
Janus h4
Janus h5
B)
0001a
01111
02338
02397
02893
Alignment Identity (%)
100
95
90
85
80
0
10k
20k
30k
C)
Alignment Identity (%)
90
80
70
Gene category
Core
Shared Accessory
Subcluster Accessory
0.0
0.1
0.2
0.3
0
5 000
10 000
15 000
20 000
D)
Ellipse subcluster
Diamond subcluster
E)
04011
F21572e04990
B5461s04202
CEA10e00035
01424
00606
Alignment Identity (%)
90
80
70
0
100k
200k
300k
