## Supplementary material for "StarCrew: A pipeline for exploratory analysis of *Starship* cargo": Supplemetary_information_file: FigureS7.pdf

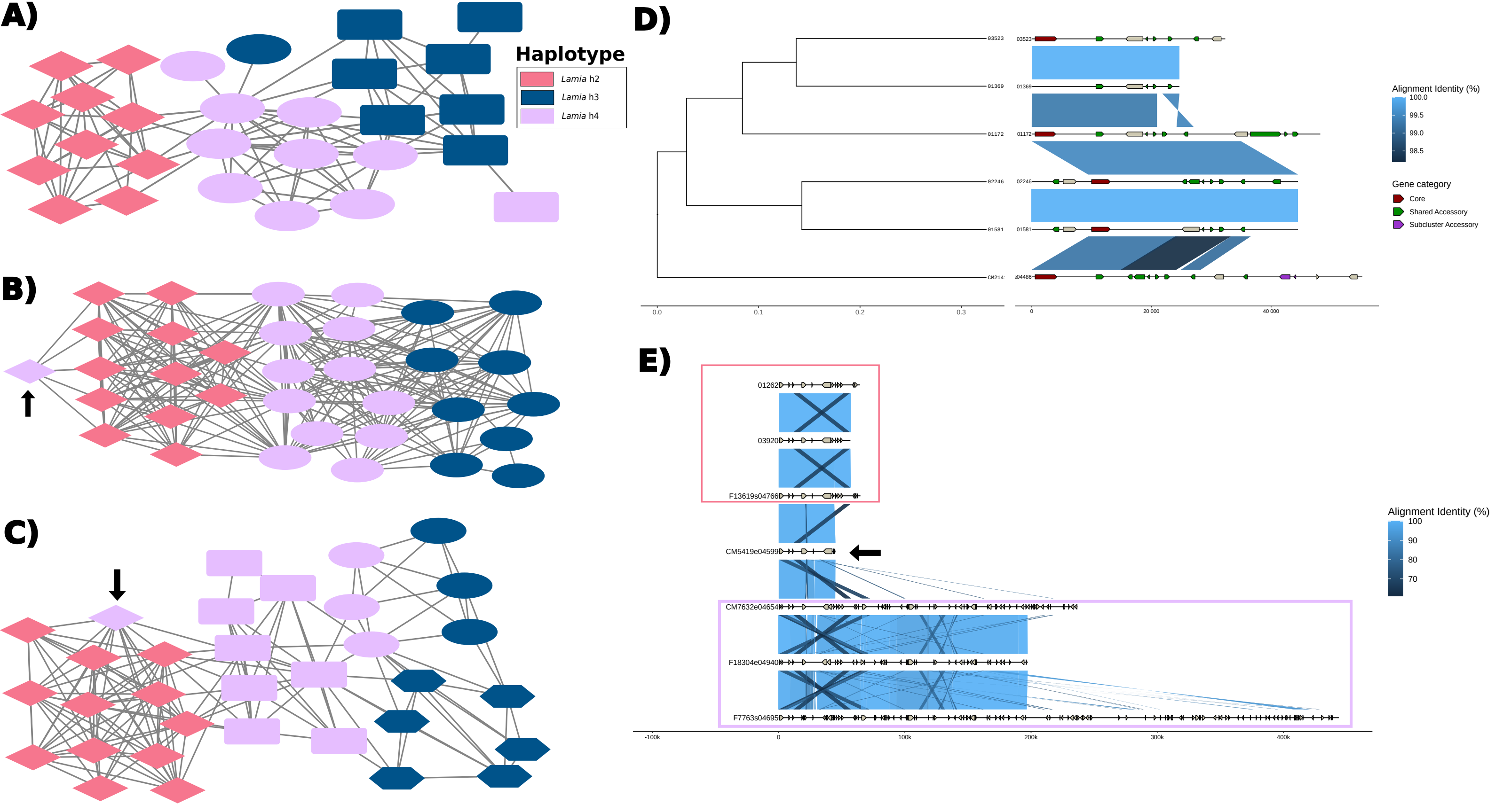
A)
Haplotype
Lamia h2
Lamia h3
Lamia h4
B)
C)
D)
03523
01369
01172
02246
01581
CM214
0.0
0.1
0.2
0.3
0
20 000
40 000
Alignment Identity (%)
100.0
99.5
99.0
98.5
Gene category
Core
Shared Accessory
Subcluster Accessory
E)
01262
03920
F13619s04766
CM5419e04599
CM7632e04654
F18304e04940
F7763s04695
-100k
0
100k
200k
300k
400k
Alignment Identity (%)
100
90
80
70
