## Supplementary material for "StarCrew: A pipeline for exploratory analysis of *Starship* cargo": Supplemetary_information_file: FigureS9.pdf

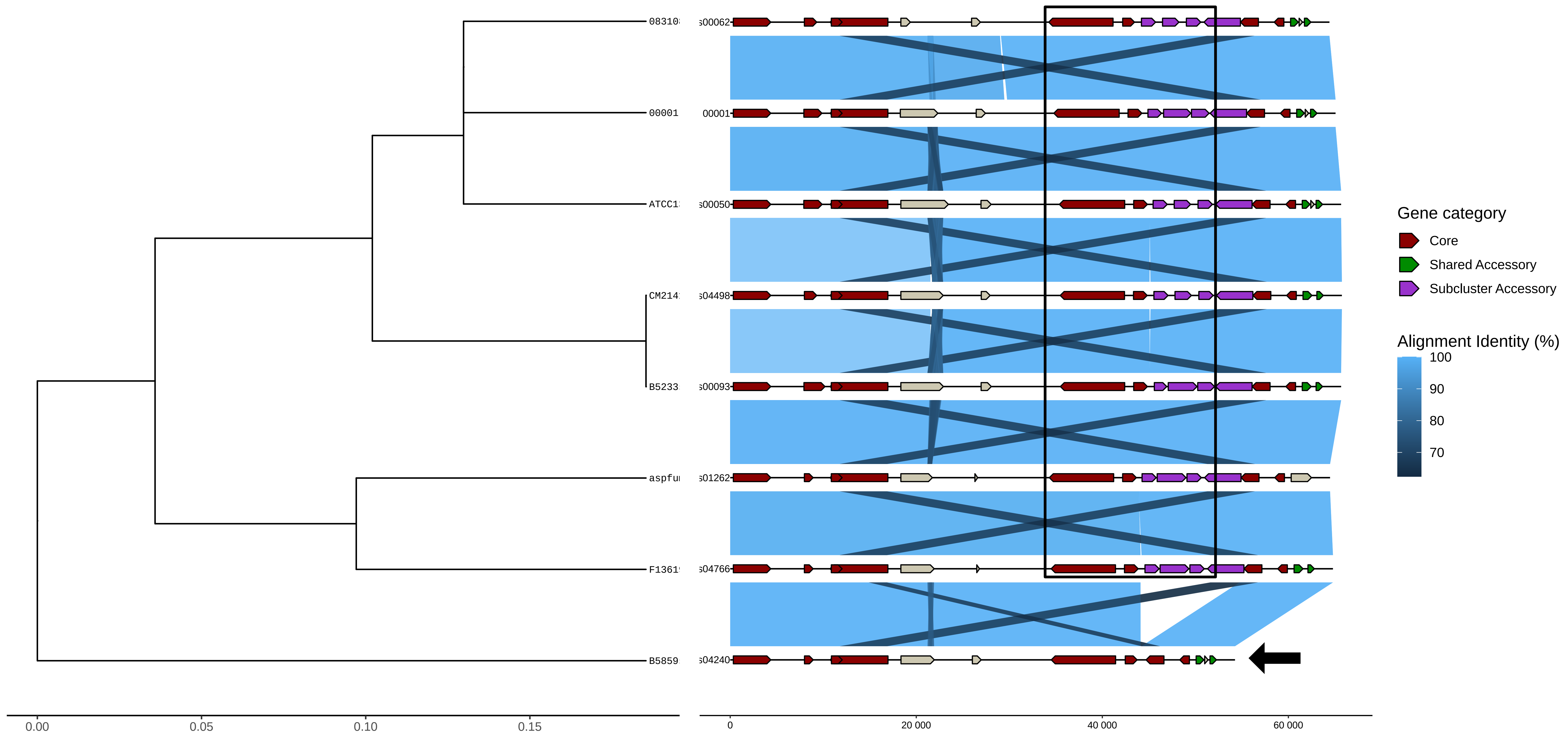

08310
00001
ATCC1
CM214
B5233
aspfu
F1361
B5859
0.00
0.05
0.10
0.15
s00062
00001
s00050
s04498
s00093
s01262
s04766
s04240
0
20 000
40 000
60 000
Gene category
Core
Shared Accessory
Subcluster Accessory
Alignment Identity (%)
100
90
80
70
