## Supplementary material for "StarCrew: A pipeline for exploratory analysis of *Starship* cargo": Supplemetary_information_file: FigureS11.pdf

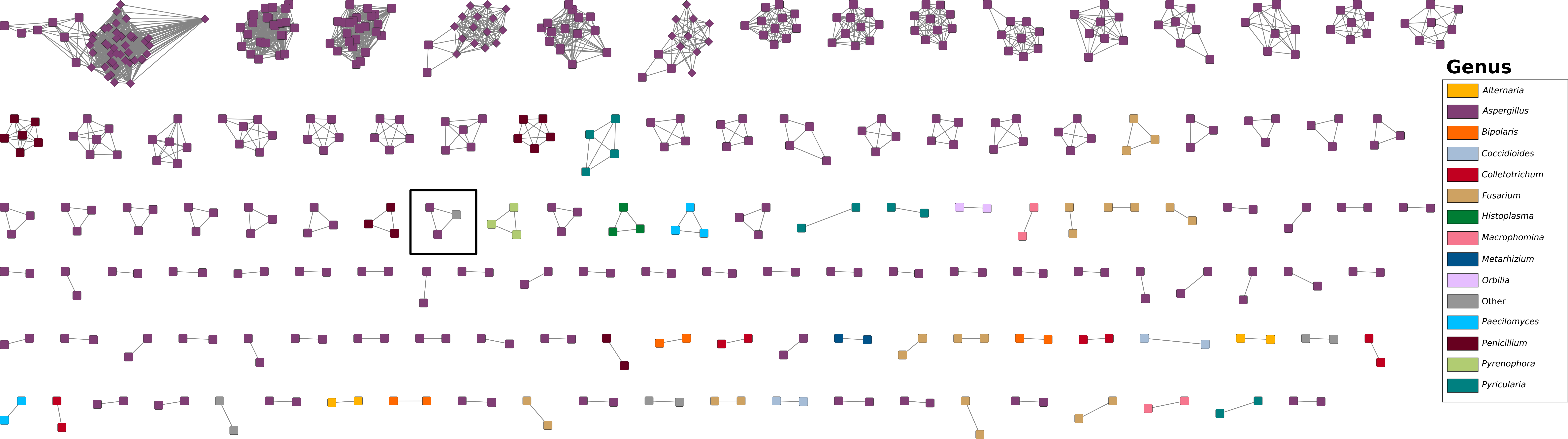
Genus
Alternaria
Aspergillus
Bipolaris
Coccidioides
Colletotrichum
Fusarium
Histoplasma
Macrophomina
Metarhizium
Orbilia
Other
Paecilomyces
Penicillium
Pyrenophora
Pyricularia
