## Supplementary material for "StarCrew: A pipeline for exploratory analysis of *Starship* cargo": Supplemetary_information_file: FigureS13.pdf

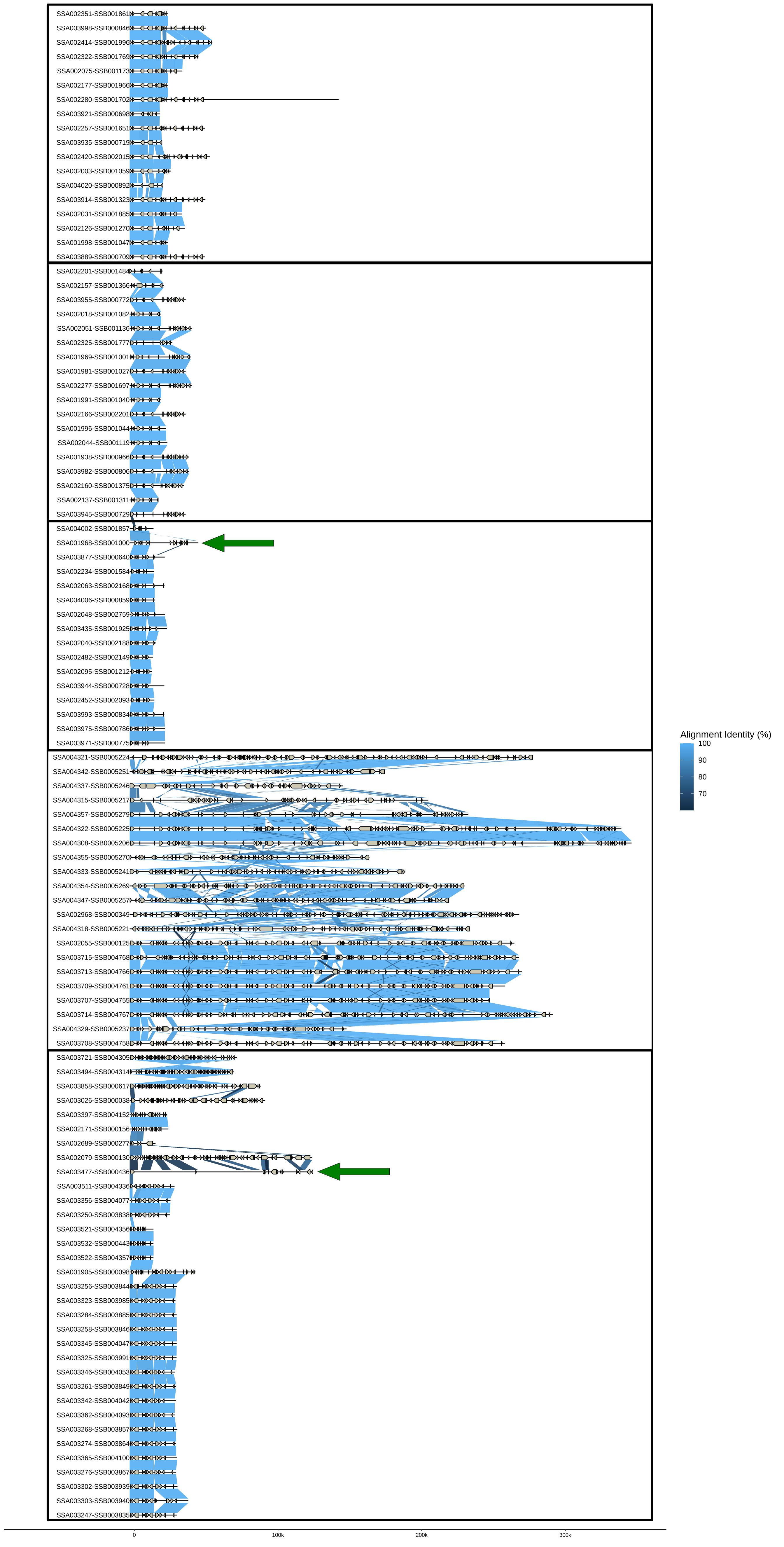
SSA002351-SSB001861
SSA003998-SSB000846
SSA002414-SSB001996
SSA002322-SSB001769
SSA002075-SSB001173
SSA002177-SSB001966
SSA002280-SSB001702
SSA003921-SSB000698
SSA002257-SSB001651
SSA003935-SSB000719
SSA002420-SSB002015
SSA002003-SSB001059
SSA004020-SSB000892
SSA003914-SSB001323
SSA002031-SSB001885
SSA002126-SSB001270
SSA001998-SSB001047
SSA003889-SSB000709
SSA002201-SSB001484
SSA002157-SSB001366
SSA003955-SSB000772
SSA002018-SSB001082
SSA002051-SSB001136
SSA002325-SSB001777
SSA001969-SSB001001
SSA001981-SSB001027
SSA002277-SSB001697
SSA001991-SSB001040
SSA002166-SSB002201
SSA001996-SSB001044
SSA002044-SSB001119
SSA001938-SSB000966
SSA003982-SSB000806
SSA002160-SSB001375
SSA002137-SSB001311
SSA003945-SSB000729
SSA004002-SSB001857
SSA001968-SSB001000
SSA003877-SSB000640
SSA002234-SSB001584
SSA002063-SSB002168
SSA004006-SSB000859
SSA002048-SSB002759
SSA003435-SSB001925
SSA002040-SSB002188
SSA002482-SSB002149
SSA002095-SSB001212
SSA003944-SSB000728
SSA002452-SSB002093
SSA003993-SSB000834
SSA003975-SSB000786
SSA003971-SSB000775
SSA004321-SSB0005224
SSA004342-SSB0005251
SSA004337-SSB0005246
SSA004315-SSB0005217
SSA004357-SSB0005279
SSA004322-SSB0005225
SSA004308-SSB0005206
SSA004355-SSB0005270
SSA004333-SSB0005241
SSA004354-SSB0005269
SSA004347-SSB0005257
SSA002968-SSB000349
SSA004318-SSB0005221
SSA002055-SSB000125
SSA003715-SSB004768
SSA003713-SSB004766
SSA003709-SSB004761
SSA003707-SSB004755
SSA003714-SSB004767
SSA004329-SSB0005237
SSA003708-SSB004758
SSA003721-SSB004305
SSA003494-SSB004314
SSA003858-SSB000617
SSA003026-SSB000038
SSA003397-SSB004152
SSA002171-SSB000156
SSA002689-SSB000277
SSA002079-SSB000130
SSA003477-SSB000436
SSA003511-SSB004336
SSA003356-SSB004077
SSA003250-SSB003838
SSA003521-SSB004356
SSA003532-SSB000443
SSA003522-SSB004357
SSA001905-SSB000098
SSA003256-SSB003844
SSA003323-SSB003985
SSA003284-SSB003885
SSA003258-SSB003846
SSA003345-SSB004047
SSA003325-SSB003991
SSA003346-SSB004053
SSA003261-SSB003849
SSA003342-SSB004042
SSA003362-SSB004093
SSA003268-SSB003857
SSA003274-SSB003864
SSA003365-SSB004100
SSA003276-SSB003867
SSA003302-SSB003939
SSA003303-SSB003940
SSA003247-SSB003835
0
100k
200k
300k
Alignment Identity (%)
100
90
80
70
