## Supplementary material for "StarCrew: A pipeline for exploratory analysis of *Starship* cargo": Supplemetary_information_file: FigureS14.pdf

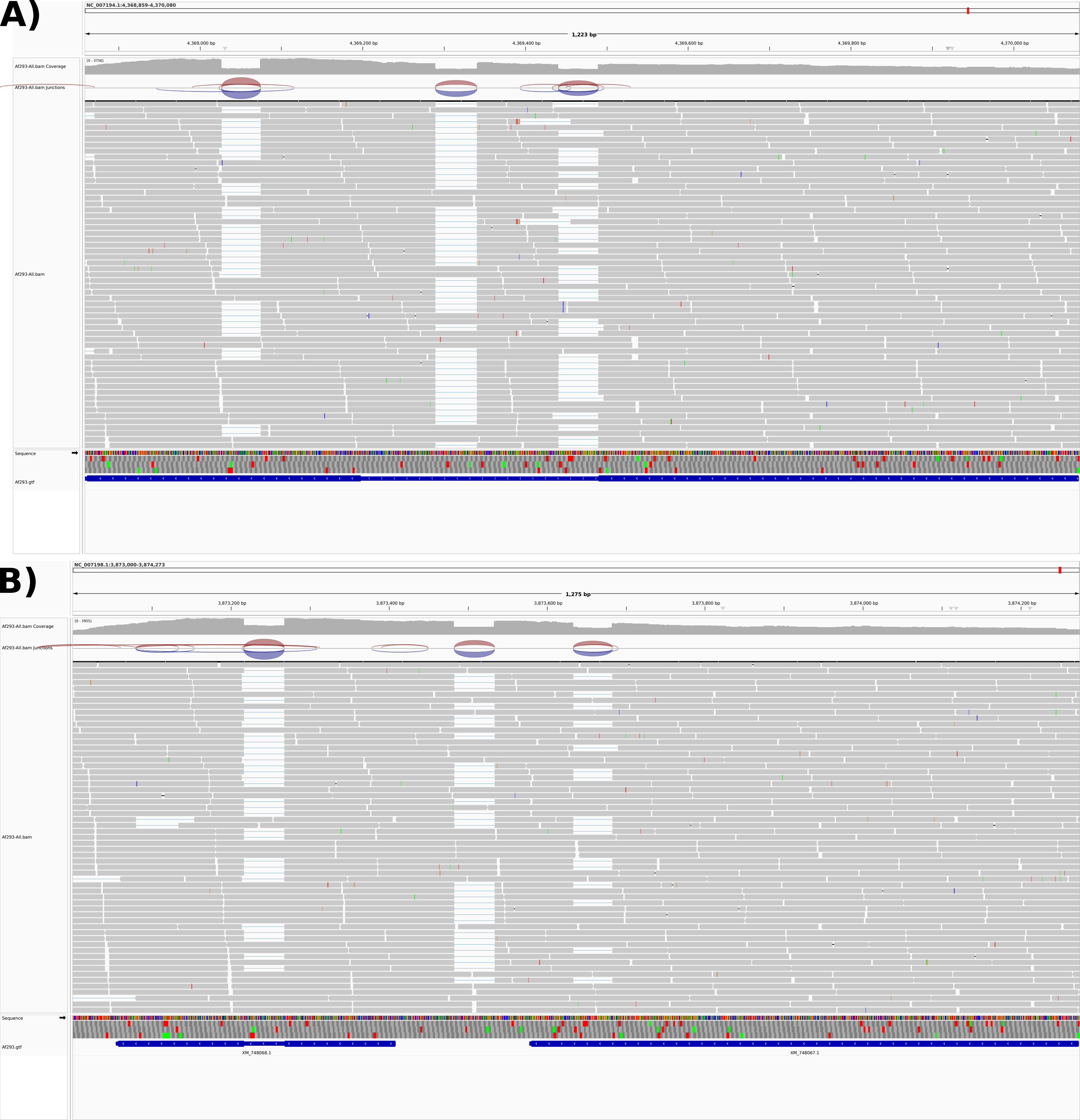
A)
NC_007194.1:4,368,859-4,370,080
1,223 bp
4,369,000 bp
4,369,200 bp
4,369,400 bp
4,369,600 bp
4,369,800 bp
4,370,000 bp
[0 - 3756]
Af293-All.bam Coverage
Af293-All.bam Junctions
Af293-All.bam
Sequence
Af293.gtf
B)
NC_007198.1:3,873,000-3,874,273
1,275 bp
3,873,200 bp
3,873,400 bp
3,873,600 bp
3,873,800 bp
3,874,000 bp
3,874,200 bp
[0 - 3955]
Af293-All.bam Coverage
Af293-All.bam Junctions
Af293-All.bam
Sequence
Af293.gtf
XM_748068.1
XM_748067.1
