## Supplementary figures and images for "StarCrew: A pipeline for exploratory analysis of *Starship* cargo"

### FigureS8.pdf

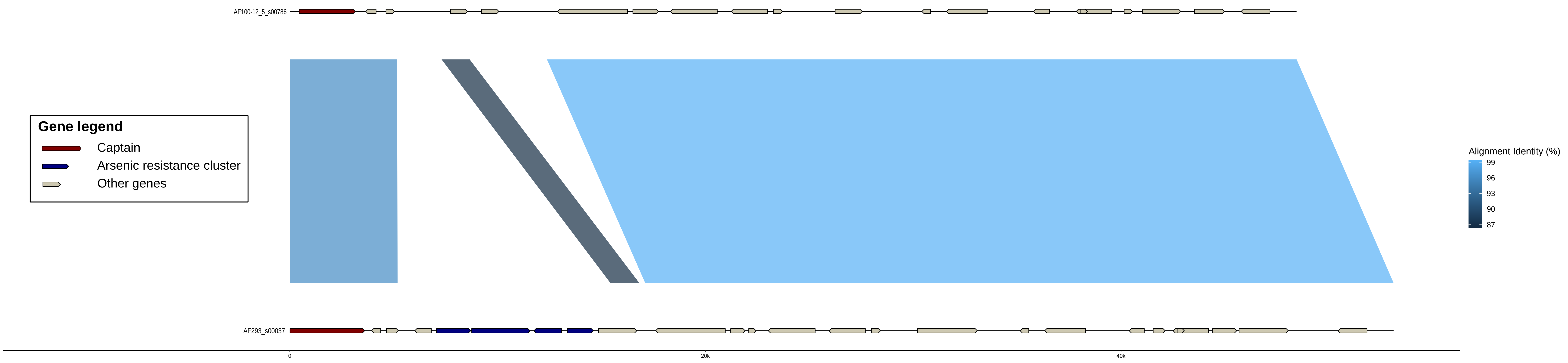

AF100-12_5_s00786
AF293_s00037
Gene legend
Captain
Arsenic resistance cluster
Other genes
Alignment Identity (%)
99
96
93
90
87
0
20k
40k

### FigureS10.pdf

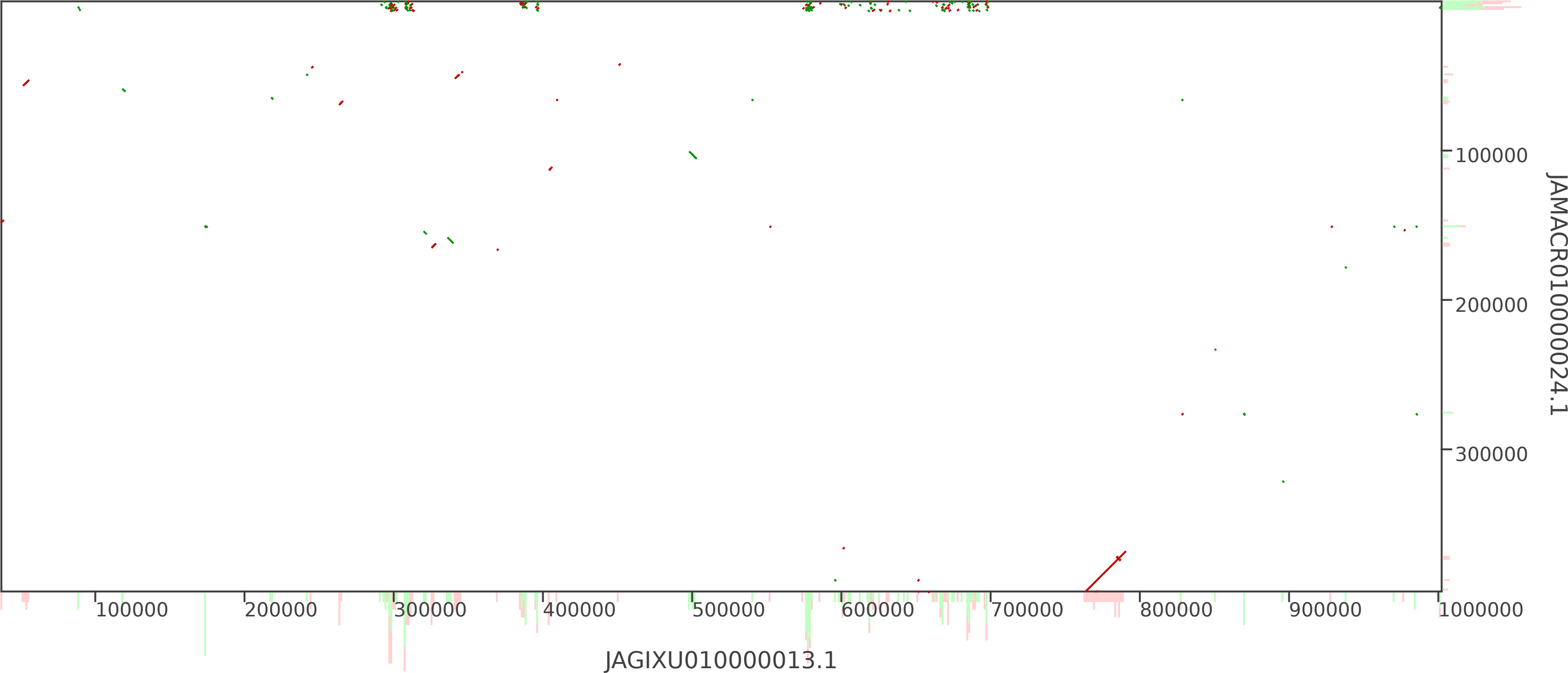

100000
200000
300000
JAMACR010000024.1
100000
200000
300000
400000
500000
600000
700000
800000
900000
1000000
JAGIXU010000013.1

### FigureS12.pdf

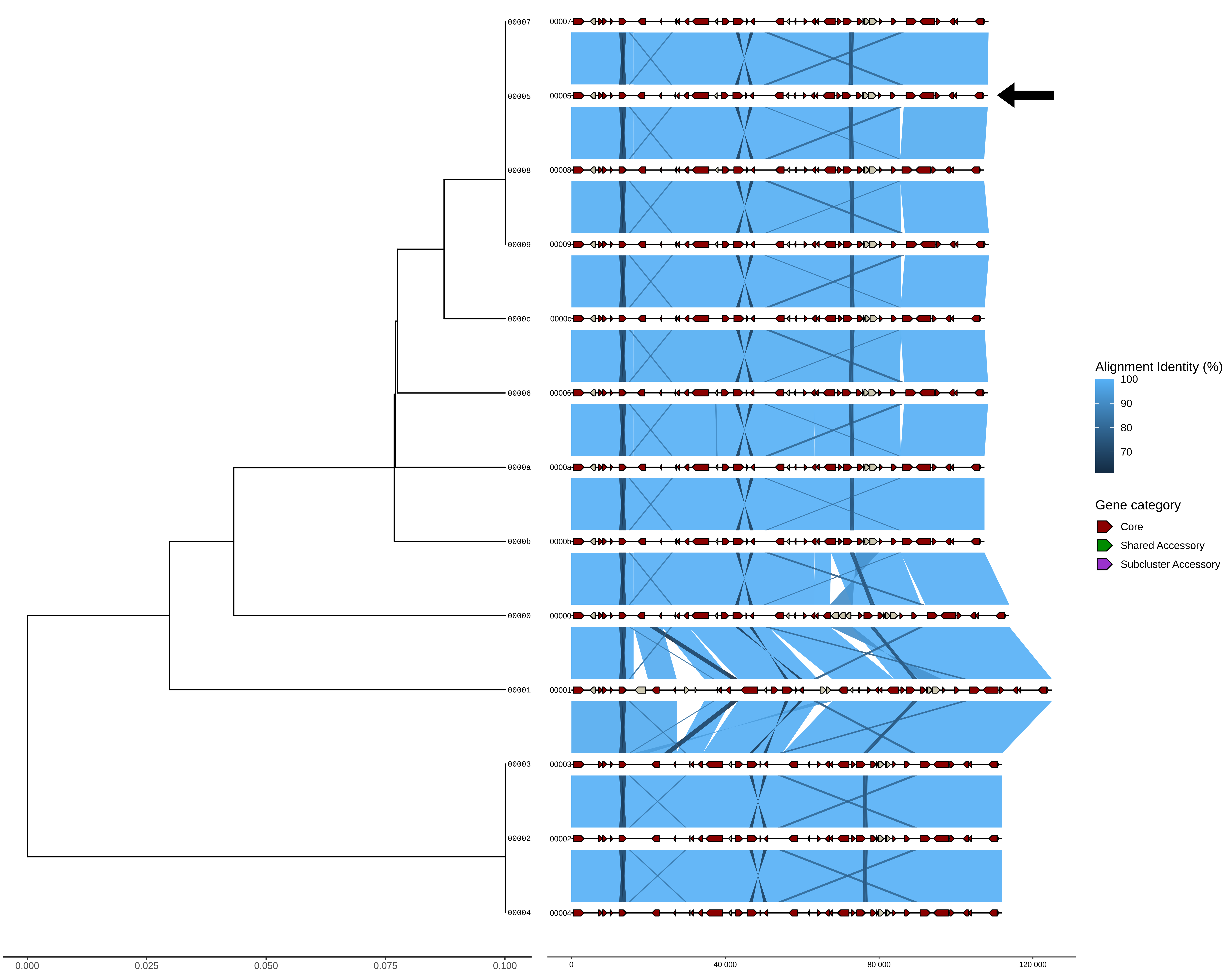

00007
00005
00008
00009
0000c
00006
0000a
0000b
00000
00001
00003
00002
00004
0.000
0.025
0.050
0.075
0.100
0
40 000
80 000
120 000
Alignment Identity (%)
100
90
80
70
Gene category
Core
Shared Accessory
Subcluster Accessory

### FigureS15.pdf

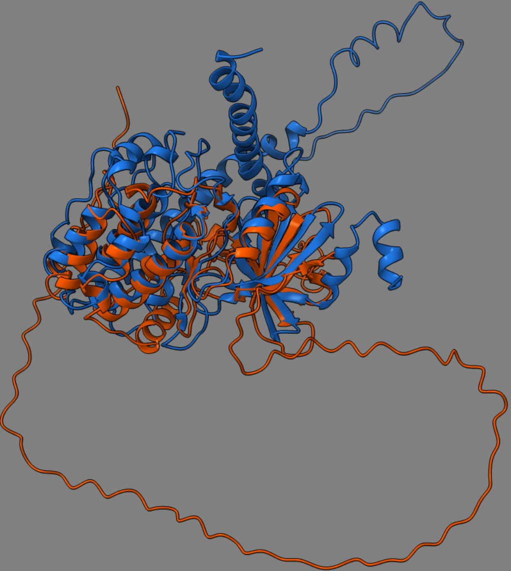
